# Mapping the complex transcriptional landscape of the phytopathogenic bacterium *Dickeya dadantii*

**DOI:** 10.1101/2020.09.30.320440

**Authors:** Raphaël Forquet, Xuejiao Jiang, William Nasser, Florence Hommais, Sylvie Reverchon, Sam Meyer

## Abstract

*Dickeya dadantii* is a phytopathogenic bacterium that causes soft rot in a wide range of plant hosts worldwide, and a model organism for studying virulence gene regulation. The present study provides a comprehensive and annotated transcriptomic map of *D. dadantii* obtained by a computational method combining five independent transcriptomic datasets: (i) paired-end RNA-seq data for a precise reconstruction of the RNA landscape; (ii) DNA microarray data providing transcriptional responses to a broad variety of environmental conditions; (iii) long-read Nanopore native RNA-seq data for isoform-level transcriptome validation and determination of transcription termination sites; (iv)dRNA-seq data forthe precise mapping of transcription start sites; (v) *in planta* DNA microarray data for a comparison of gene expression profiles between *in vitro* experiments and the early stages of plant infection. Our results show that transcription units sometimes coincide with predicted operons but are generally longer, most of them comprising internal promoters and terminators that generate alternative transcripts of variable gene composition. We characterise the occurrence of transcriptional read-through at terminators, which might play a basal regulation role and explain the extent of transcription beyond the scale of operons. We finally highlight the presence of noncontiguous operons and excludons in the *D. dadantii* genome, novel genomic arrangements that might contribute to the basal coordination of transcription. The highlighted transcriptional organisation may allow *D. dadantii* to finely adjust its gene expression programme for a rapid adaptation to fast changing environments.

**IMPORTANCE:** This is the first transcriptomic map of a *Dickeya* species. It may therefore significantly contribute to further progress in the field of phytopathogenicity. It is also one of the first reported applications of long-read Nanopore native RNA-seq in prokaryotes. Our findings yield insights into basal rules of coordination of transcription that might be valid for other bacteria, and may raise interest in the field of microbiology in general. In particular, we demonstrate that gene expression is coordinated at the scale of transcription units rather than operons, which are larger functional genomic units capable of generating transcripts with variable gene composition for a fine-tuning of gene expression in response to environmental changes. In line with recent studies, our findings indicate that the canonical operon model is insufficient to explain the complexity of bacterial transcriptomes.

## INTRODUCTION

Classically, bacterial transcription is described with the model of Jacob and Monod based on operons, defined as sets of contiguous and functionally-related genes co-transcribed from a single promoter up to a single terminator (1). In recent years however, accumulating studies demonstrated that most operons actually comprise internal promoters and terminators, generating transcripts of variable gene composition, generally in a condition-dependent manner (2, 3, 4, 5). This phenomenon, also known as suboperonic regulation (6), might be compared to alternative splicing in eukaryotes (7) and demonstrates a higher complexity of bacterial transcriptional landscapes than previously thought. Besides, transcription has been shown to extend beyond operons (3, 8), the latter being actually part of larger functional genomic units, referred to as transcription units (TUs) throughout the manuscript.

While transcriptomic maps have been established for various bacteria including *Escherichia coli* (9), *Salmonella enterica* (10), *Bacillus subtilis* (2), *Streptococcus pneumoniae* (4), *Campylobacter jejuni* (11), *Clostridium beijerinckii* (12), *Mycobacterium tuberculosis* (13), *Mycoplasma pneumoniae* (14), and the phytopathogen *Xanthomonas campestris* (15, 16), they are still lacking for *Dickeya species*. This study aims to provide the first comprehensive and annotated transcriptomic map of *Dickeya dadantii*, a Gram-negative phytopathogenic bacterium representative of the *Dickeya* genus that causes soft rot, a severe disease leading to tissue maceration and eventually plant death (17) in a wide range of plant hosts worldwide, including agriculturally important crops (18, 19, 20, 21, 22).

The infection process involves an asymptomatic phase, where bacteria remain latent, penetrate and colonise plant tissues, consuming simple sugars and small soluble oligosaccharides available in the plant apoplast to grow exponentially (23). In this compartment, bacteria are exposed to acidic conditions (24) and oxidative stress (25) resulting from plant defences. When all nutrients are consumed in the apoplast, the symptomatic phase initiates. Bacteria produce plant cell wall degrading enzymes (mainly pectinases) leading to the soft rot symptoms, and start cleaving pectin, which is used as a secondary carbon source for a new round of growth (26). By causing a total destruction of plant cells, the maceration of plant tissues releases both vacuolar and cytoplasmic components in the apoplast, exposing the bacteria to osmotic stress (23).

In order to characterise the *D. dadantii* transcriptional landscape, we used a combination of transcriptomic data generated *in vitro* in a broad range of growth and stress conditions reflecting some of the key environmental signals encountered during the plant infection process, and ensuring optimal reproducibility and quality of analysed RNAs (27, 28). Different techniques were used, providing complementary knowledge: high-resolution Illumina paired-end RNA-seq; DNA microarray; Nanopore native RNA-seq; dRNA-seq. These data were combined using an integrative computational method developed for this study, allowing the inference of the RNA landscape and a validation of co-expression occurring among genes of the same TU. This analysis provides a detailed and annotated map of the TUs defining the *D. dadantii* transcriptome, *i.e.*, the sets of contiguous co-expressed genes. We then quantitatively map transcription start and termination sites in the investigated conditions, and analyse the associated predicted promoter and terminator motifs. We show that TUs sometimes coincide with predicted operons but are generally longer, most of them exhibiting internal promoters and terminators. We characterise the occurrence of transcriptional read-through at terminators, a mechanism proposed as a basal coordinator and regulator of gene expression yet never explored in phytopathogens and still poorly understood across genomes in general. We finally detect putative noncontiguous oper-ons and excludons in the *D. dadantii* genome. In order to validate the obtained transcriptional map, we analyse available *in planta* expression data, and show that TUs inferred from *in vitro* cultures are also co-expressed during the early stages of plant infection (29), suggesting that many of the analysed features are used by *D. dadantii* in the pathogenic context. This transcriptomic map might serve as a community resource to help elucidating the regulation of *D. dadantii* gene expression, including its virulence programme. It also provides insights into basal rules of coordination of transcription that might be valid for other bacteria, specifically for other *Dickeya* species for which a core genome of 1300 genes was identified by comparative genomics (30).

## RESULTS AND DISCUSSION

### Characterisation of *Dickeya dadantii* transcription units

In order to generate a biologically relevant transcriptional map of *D. dadantii,* we combined and integrated four sets of transcriptomic data obtained from *in vitro* cultures subjected to different sugar sources, environmental stress factors (acidic, oxidative, osmotic stress), and variations of DNA supercoiling, reflecting a variety of conditions also encountered by bacteria in the course of plant infection. A fifth set obtained from bacteria grown *in planta* was used for validation. These data were collected by different experimental methods providing complementary information, as follows (a more detailed description of the datasets is provided in Materials and Methods).

Dataset 1 was generated from high-resolution Illumina paired-end, strand-specific RNA-seq covering 6 growth conditions: M63 minimal medium supplemented with sucrose, addition of polygalacturonate (PGA), a pectic polymer present in plant cell wall (31), and treatment by novobiocin, which induces a global and transient chromosomal DNA relaxation (32) in exponential or in early stationary phase. By providing short but precise sequencing reads at single base-pair resolution and high sequencing depth, this dataset yields precise and quantitative information on the RNA landscape.

Dataset 2 was generated from DNA microarray data covering 32 growth conditions, involving the presence of PGA and leaf extracts, and in each medium, a separate exposure to acidic, oxidative or osmotic stresses (28). This dataset provides a quantitative catalogue of genes’ responses to a more comprehensive and detailed range of conditions than dataset 1, albeit of weaker spatial resolution.

Dataset 3 was generated from long-read Nanopore native RNA-seq in M63 minimal medium supplemented with glucose and PGA, pooled from samples obtained in both exponential and early stationary phases. This method allows native RNAs to be sequenced directly as near full-length transcripts from the 3’ to 5’ direction, with a weaker depth than the previous datasets. Only a few transcriptomes were analysed by this technique, mostly from viral and eukaryotic organisms (33, 34, 35, 36), and, to our knowledge, a single prokaryotic one (37). This dataset provides a direct isoform-level validation of the TUs, and an accurate definition of transcription termination sites.

Dataset 4 was generated from differential RNA sequencing (dRNA-seq) experiments carried out on four samples obtained by pooling RNAs from the large variety of environmental conditions of dataset 2 followed by treatment with Terminator exonuclease (TEX) prior to sequencing. TEX enzyme degrades processed 5’-monophosphate RNAs and consequently enriches the samples in primary 5’-triphosphate end transcripts (38), thus locating transcription start sites at single-nucleotide resolution.

Finally, dataset 5 was generated from *in planta* DNA microarray data, 6 and 24 hours post-inoculation of the model plant *Arabidopsis thaliana* (29), during the early stages of infection. Bacterial RNAs are difficult to isolate from plant tissues, especially during the symptomatic phase where phenolic compounds accumulate in decaying tissues, explaining the lack of transcriptomic data during the late stages of infection. In spite of a limited variety of conditions, this dataset allows a comparison of gene expression profiles between *in vitro* and *in planta* experiments, and was used to validate the level of co-expression of genes within TUs during the early stages of plant infection.

This collection of diverse and complementary transcriptomic datasets provided a solid ground for precisely characterising the *D. dadantii* transcription units, rather than basing our analysis on genomic data alone as in most operon predictors (intergenic distances between genes, functional links among products). The employed algorithm is described in details in Materials and Methods. Shortly, in a first step, we analysed the RNA landscape from Illumina paired-end strand-specific RNA-Seq (dataset 1), ensuring good resolution and sufficient sequencing depth to obtain a quantitative signal for all genes. These data also allowed us to uncover 50 putative coding genes previously unannotated, most of which exhibiting sequence homology with proteins from the *Dickeya* genus (Supplementary Tab. S1D). Putative TUs were defined by fusing adjacent genes as long as RNA fragments were found in their intergenic region, a signature of co-transcription. Secondly, if genes within the same putative TU are indeed co-transcribed, they should exhibit strong correlation of expression in a wider range of conditions than those of dataset 1. This analysis was carried using the diversity of samples in our DNA microarray data (dataset 2), based on a customised hierarchical clustering framework (39). This second criterion (correlation of expression) provided an orthogonal cross-validation compared to the first one (intergenic RNA signal), and yielded a total of 2028 putative TUs along the *D. dadantii* genome. In a third step, these TUs were validated based on Nanopore native RNA-seq (dataset 3). We tested the presence of long native RNA reads overlapping adjacent genes belonging to the same TU, thus yielding a direct evidence of co-transcription. For 16% of adjacent gene pairs, no conclusion could be drawn because of insufficient coverage. For the others, co-transcription was confirmed in 92% of the cases; for the remaining 8%, the absence of a common RNA might be indicative of false positives, but for some of them, may also be due to the weak number of culture conditions included in dataset 3. Since the large majority of TUs defined from datasets 1 and 2 match the observations of Nanopore native RNA-seq, we favoured the latter hypothesis and retained all of them, with a confidence level reflecting the presence or absence of overlapping RNA reads (Supplementary Tab. S1A).

With this approach, we mapped the first layer of transcription organisation in *D. dadantii*. According to our findings, the 4211 protein-coding genes are organised into 2028 transcription units (provided in Supplementary Tab. S1A), among which 1118 are monocistronic and 910 are polycistronic, ranging from 2 to 28 genes (Fig. 1A, 1B and Supplementary Tab. S5). At the genomic scale, we compared our results with those of Rockhopper, a popular operon predictor that uses expression data as well as genomic information as input (40). 45% of predicted operons exactly coincide with a TU in our analysis (Fig. 1D), including known examples such as *smtAmukFEB* involved in chromosome partitioning (Fig. 2A) (41). Besides, many identified TUs are likely operons of unknown functions and features (Fig. 2B), which represent interesting starting points to discover new transcriptional functional units. Remarkably, TUs are generally longer than predicted operons: the average TU (including monocistronic ones) contains 2.1 genes and the average polycistronic TU contains 3.4 genes, against 1.6 and 3.1 respectively for predicted operons (Fig. 1C). Almost three quarters (73.5%) of all genes are co-transcribed in TUs, against 56.9% for predicted operons (Fig. 1A, Supplementary Tab. S5). Our results indicate that TUs are indeed larger functional genomic units, since 45% of predicted operons are extended by at least one gene (Fig. 1D), in agreement with recent findings in *E. coli* based on long-read sequencing (3).

**FIG 1.**
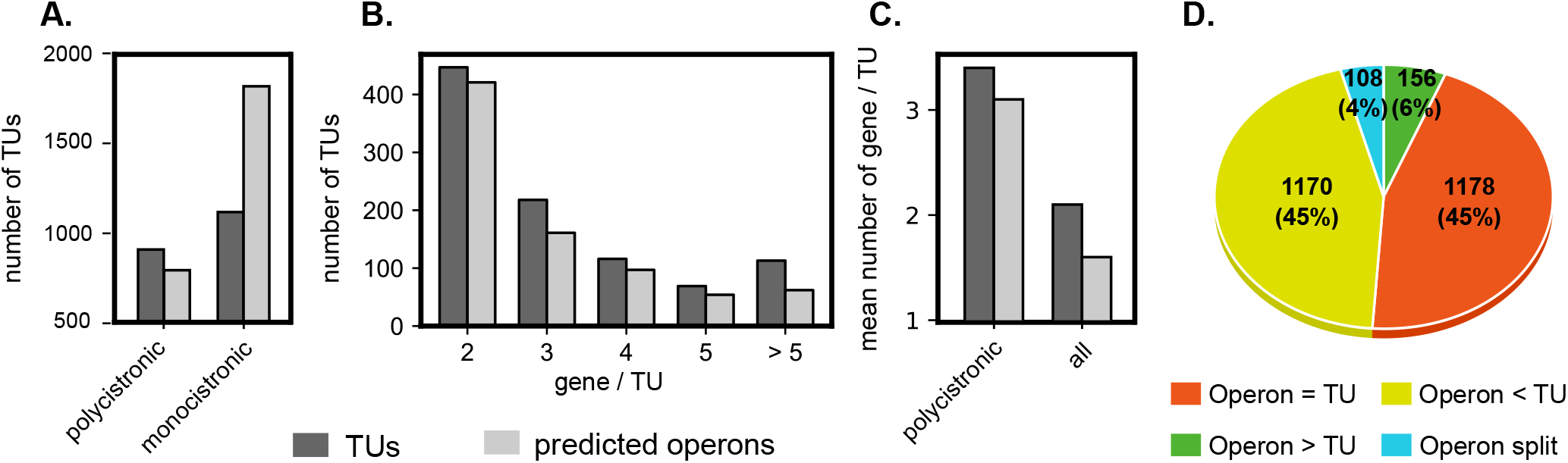
(A) Repartition of monocistronic and polycistronic TUs identified by our analysis and comparison to predicted operons. (B) Size distributions. (C) Average number of genes per TU. (D) Fate of predicted operons that are mostly found as or within TUs in our algorithm.

**FIG 2.**
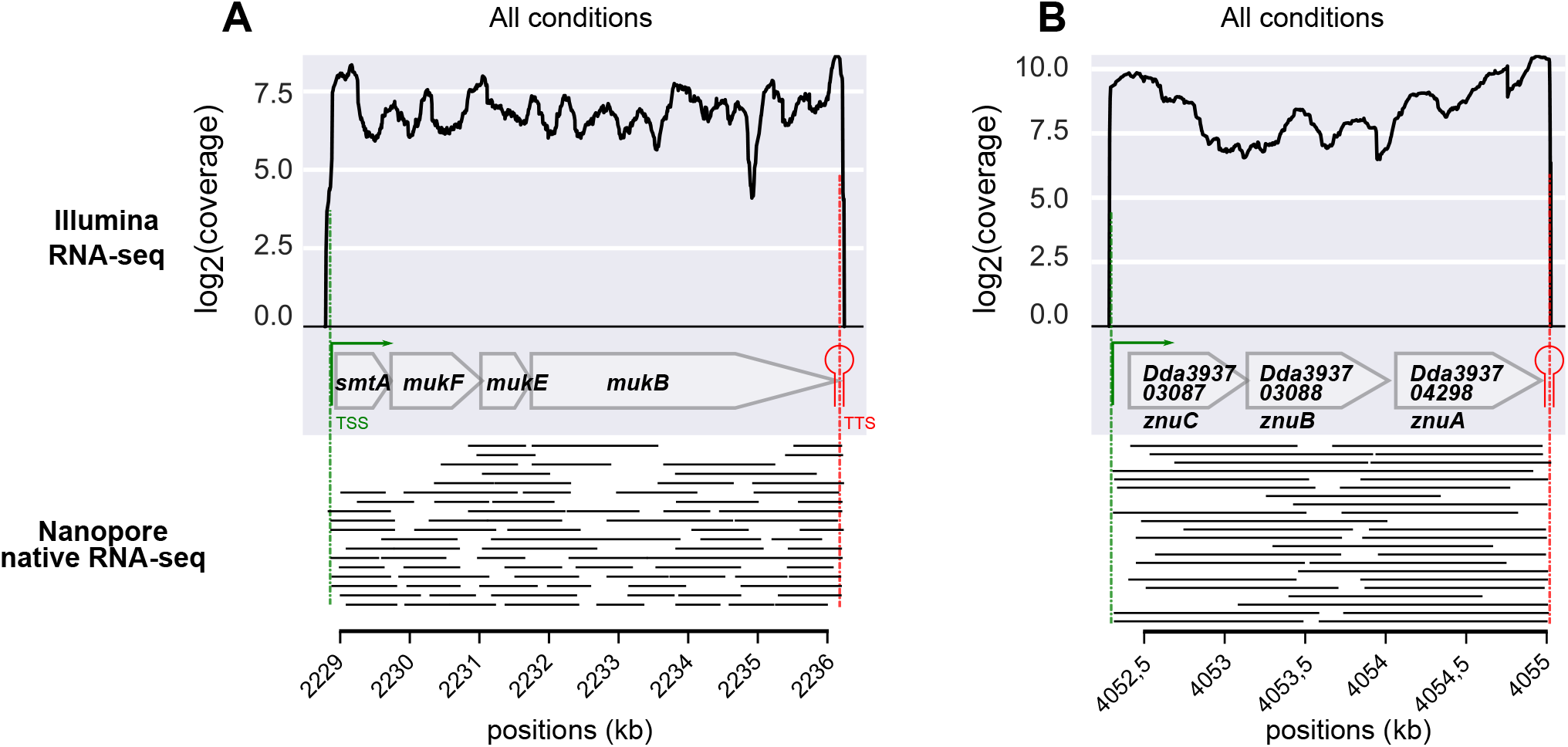
Transcription units identified by our approach, and coinciding with operons. (A) Example of a known operon (*smtAmukFEB*). The bottom panel shows the coordinates of the long native RNA reads sequenced by Nanopore. (B) Identification of a new TU exhibiting uniform read coverage and strong internal cross-correlations (Supplementary Fig. S1A), clearly indicative of an operon. Its function was unknown but a homology analysis revealed that it corresponds to the cluster of genes *znuCBA*, a *Zn*^2+^ uptake system (47). Long reads are observed for all adjacent gene pairs in Nanopore native RNA-seq data, and even a fragment carrying the three genes for *znuCBA*.

As an example, the *sapABCDF* operon encoding a transporter involved in antimicrobial peptide resistance and virulence in numerous bacteria including *D. dadantii* (42) is extended to include the enoyl-acyl carrier protein reductase *fabI* that catalyses an essential step in the biosynthesis of fatty acids of the membrane (43) (Fig. 3B). It might be noted that *fabI* has a different genomic location in *E. coli* and is consequently not co-transcribed with *sapABCDF* in that species (44) although this synteny is conserved in other *Dickeya* genomes, showing that TUs can merge and/or vary over time at the evolutionary scale. Since these genes are functionally unrelated (except for a general relation with the membrane), the biological relevance and putative role of this event requires further investigation.

**FIG 3.**
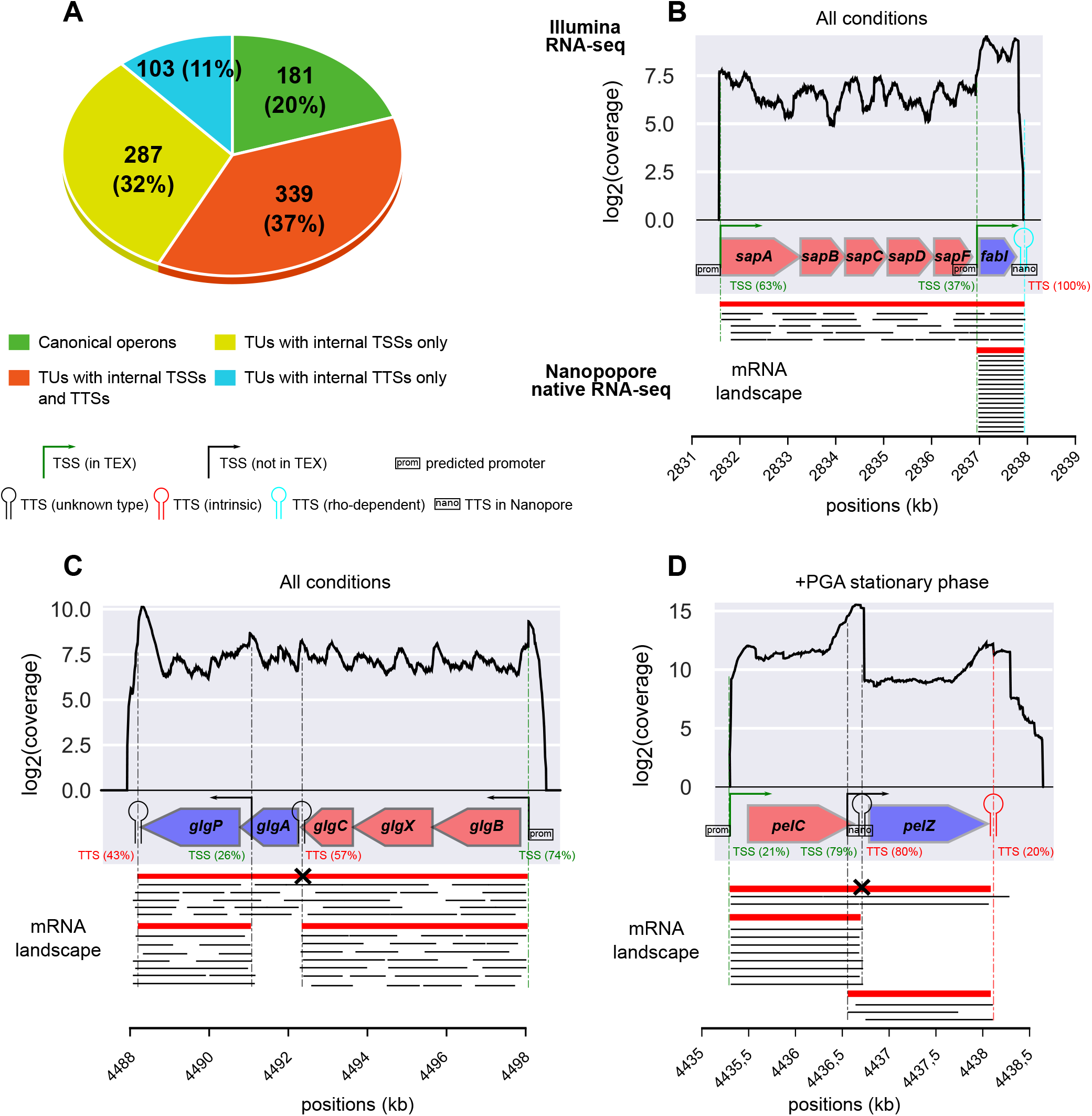
(A) Characteristics of TUs. (B) The *sapABCDF* and *fabI* genes, predicted by Rockhopper (40) as two separate (red and blue) operons, were identified as a single TU, with a strong internal TSS expressing *fabI* alone. The bottom panel indicates the different isoforms (red) and the long reads sequenced by Nanopore native RNA-seq (black). The latter overlap all adjacent gene pairs, providing a direct evidence for co-transcription. (C) The *glg* genes were identified as a single TU (involving several isoforms) containing two separate predicted operons (blue and red genes), as suggested by the uniform read coverage, long reads from Nanopore native RNA-seq (bottom), and in line with results in *E. coli* (3, 54, 46). (D) Identification of the *pelCZ* TU with different isoforms depending on the condition, as previously determined (55). The two genes are split into different operons by Rockhopper. A strong internal TSS, followed by a strong TTS, contributes to the complexity of its expression (see text). Long reads corresponding to the different mRNA isoforms (*pelC, pelZ, or pelCZ*) are observed.

The *glg* genes involved in glycogen metabolism constitute another instructive example. They were initially classified in two separate operons in *E. coli* (45), and later identified as a single TU involving alternative transcripts of variable gene composition depending on growth conditions (46). The latter is also true in *D. dadantii* according to our findings (Fig. 3C), illustrating how transcription extends beyond the scale of the operon.

### Genome-wide identification of *D. dadantii* transcription start and termination sites

Once *D. dadantii* transcription units were defined, the next step was to elaborate a map of transcription start sites (TSSs) and transcription termination sites (TTSs) for each TU along the genome. First, as mentioned above, dRNA-seq experiments were carried out to build a large library of 9313 putative TSSs at high-resolution (38) covering a wide range of of *in vitro* cultures under growth and stress conditions also encountered during plant infection (dataset 4, Supplementary Tab. S2A). These were obtained by treating the RNA samples with TEX prior to sequencing, and the TSSer workflow was applied for a precise determination of TSS positions (48), followed by visual curation (Materials and Methods). For TTSs, two sets of putative positions were generated based on (i) Nanopore native RNA-seq (dataset 3), where transcripts are sequenced from the 3’ ends, allowing the detection of 1165 TTS positions based on the enrichment of these ends downstream of gene stops (Supplementary Tab. S2D); (ii) genome-wide predictions of termination sites, based on the two main mechanisms of transcription termination in bacteria. 3564 Rho-independent (intrinsic) TTSs and 5851 Rho-dependent(regulated)TTSs(49)were predicted using ARNold (50)and RhoTermPredict (51) programmes respectively (Supplementary Tabs. S2B and S2C).

A quantitative mapping of the transcription landscape was then performed in order to estimate the contribution of each TSS/TTS to its TU. While most comparable maps define TSSs/TTSs by their position only, we exploited the complementarity of the input data to also systematically analyse their magnitude (or strength) in the investigated conditions. The +TEX libraries, Nanopore reads and TTS predictions are not suitable for the latter purpose, which required building a second list of TSSs and TTSs of poorer resolution but quantitative magnitude from the non-treated paired-end RNA-seq data (dataset 1). Briefly, TSSs and TTSs were defined based on the enrichment in RNA fragment starts and stops upstream of gene starts and downstream of gene stops respectively, and the number of fragments associated to these sites across all samples was considered as the global strength. The lists obtained with the three methods (from datasets 1, 3 and 4) were then merged into a unified list of TSSs/TTSs of optimal spatial resolution, quantitative magnitude, and with an estimated level of confidence depending on the level of agreement between these datasets (see Materials and Methods). These TSSs and TTSs were then assigned to the TUs. In order to eliminate many very weak internal TTSs/TTSs (most of which likely have poor biological relevance), the latter were retained only if they yielded at least 15% of the total start/stop magnitude of the TU and were thus used at least in some of the investigated conditions. As a result, we defined a total of 2595 TSSs and 1699 TTSs (including internal ones) over all TUs (Supplementary Tab. S1A to S1C). Inevitably, some alternate TSSs/TTSs may be absent from these lists if they are specifically used in conditions not included in our datasets. Finally, a scan for promoter motifs, conducted with bTSS-finder(52), identified promoters upstream of 1848 (71%) TSSs in total (Supplementary Tab. S1B and Fig. 3). The absence of detected promoters for the remaining 29% TSSs was expected due to the limitations of such predictors (53). To evaluate the quality of our TSS definition, we compiled all experimentally determined TSSs in *D. dadantii* (by primer extension), and compared their positions to our findings (Supplementary Tab. S3). 45% displayed exactly the same position, 38% were distant by less than 5 nucleotides, and only 17% were distant by more than 6 nucleotides. Manually-annotated promoter elements from these studies also match our findings well (Supplementary Tab. S3).

### Characterisation of a complex transcriptional landscape

The quantitative mapping of TSSs and TTSs allowed us to refine the comparison of TUs and operons presented above. According to our findings, only 20% of polycistronic TUs (181) exhibit a single promoter and terminator (Fig. 2 and 3A) and thus fit into the classical definition of operons, and only 47% of these (85) are predicted as such by Rockhopper. The 80% remaining TUs (729) are complex (Fig. 3A). 32% (287) have at least one internal TSS without any internal TTS, such as *sapABCDFfabI* (Fig. 3B). 37% (339) have both internal TSS(s) and TTS(s), such as *glgBXCAP* (Fig. 3C) and *pelCZ* (Fig. 3D). Finally, 11% (103) have at least one internal TTS without any internal TSS such as *rhlB-gppA-pehV* (Fig. 4A), *pelD-paeY-pemA* (Fig. 4B) and *gcvTHP* (Fig. 6). Most *D. dadantii* TUs can consequently generate alternative transcripts of variable gene composition, resulting in a dense and complex transcriptional landscape.

**FIG 4.**
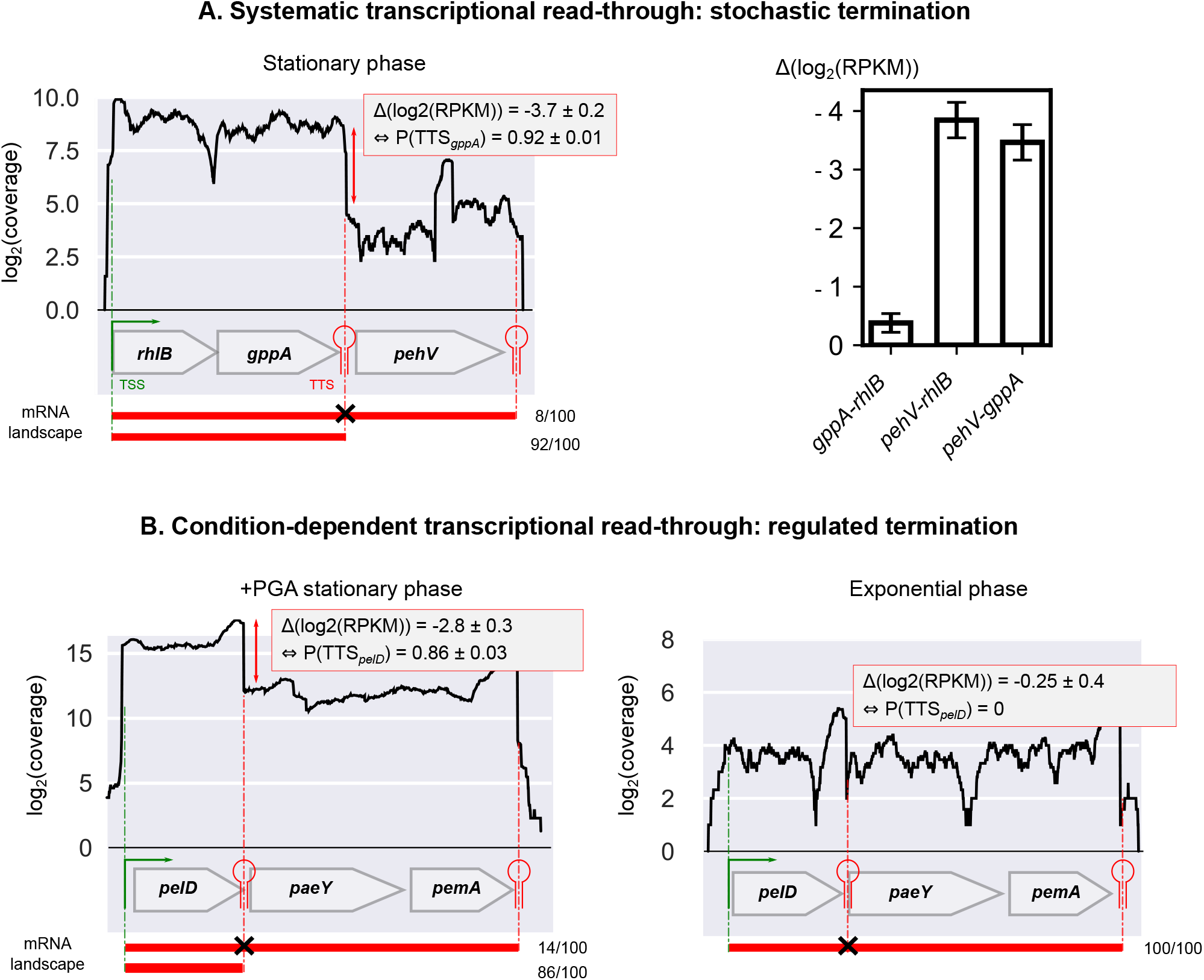
Quantification of transcriptional read-through. (A) Non-conditional read-through: example of the *rhlB-gppA-pehV* TU (left panel). The first two genes are homogeneously transcribed among conditions, resulting in an expression variation Δ(*log*_2_(*RPKM*)) close to 0 (right panel, 95% confidence intervals are shown), while the intrinsic TTS downstream of *gppA* is stochastically over-stepped in 8 ± 1% of transcripts (*P*(*TSS_gppA_*) = 0.92 ± 0.01), resulting in two different isoforms (red). (B) Condition-dependent read-through: example of the *pelD-paeY-pemA* TU. A TTS is identified downstream of *pelD* in agreement with previous studies (75). Its termination probability is regulated and depends on growth phase and presence of PGA (0.86 ± 0.03 vs 0), besides a global up-or down-regulation of the whole TU. All mRNA isoforms are observed in Nanopore native RNA-seq data (Supplementary Fig. S2A and S2B).

A notable feature of complex TUs is the heterogeneity of transcription levels along the genes due to internal TSSs / TTSs, usually in a condition-dependent manner, resulting in a moderate correlation in the expression of genes within the TU (9). As an example, in the *sapABCDFfabI* TU (Fig. 3B), *fabI* is expressed both as part of the entire transcript and as an independent transcript generated from a strong internal TSS, explaining the lower correlation between *fabI* and the remaining genes (Supplementary Fig. S1B). In *glgBXCAP* (Fig. 3C), alternative transcripts of variable gene composition can be generated depending on TSS and TTS usage. Another example relevant to plant infection is the *pelCZ* cluster (Fig. 3D) encoding two endopectate lyases secreted by *D. dadantii* which degrade pectin contained in plant cell walls (56). The substrates of Pel enzymes are pectic oligomers, *e.g.* PGA, that act as inducers of *pel* expression (31). The *pelCZ* genes were previously shown by Northern blotting to be cotranscribed into a single polycistronic transcript under inducing conditions by PGA, in addition to the two monocistronic mRNAs encoded by *pelC* or *pelZ* under non-inducing conditions (55). Our present findings are in full agreement with these observations, as *pelCZ* is detected as a single TU harbouring one internal TSS and TTS, each giving rise to monocistronic transcripts. In our data, *pelCZ* expression profiles are similar in presence or absence of PGA in spite of a drastically different global expression level (Supplementary Fig. S1D), suggesting that in absence of inducer, this very low level previously prevented a reliable detection of the entire transcript. Altogether, our findings clearly indicate that the canonical operon model is insufficient to explain the complexity ofthe *D. dadantii* transcriptional landscape, in line with results in many other organisms (2, 3, 4, 5). The existence of alternative entry and exit points for RNA Polymerase inside TUs allows the cells to adjust the relative expression level of adjacent genes within a global coordination of expression of the entire TU (Fig. 3) that may allow, in the case of *D. dadantii* during plant infection, a rapid adaptation to changing environment.

### Transcriptional read-through, the root of transcription extension?

We showed that transcription units comprise predicted operons, yet are generally longer. This extension of transcription might, in part, result from the ability of RNA Polymerase to stochastically override an imperfect terminator by a mechanism referred to as transcriptional read-through (3, 8). The latter has long been identified in specific operons (57, 58, 59) and was shown more recently to be widespread in bacterial genomes (2, 3, 8), where it may in fact play a basal coordination and regulation role (5). A condition-independent rate of stochastic termination might result in the co-expression of the genes located before and after the TTS (as in a classical operon), but with a reduced transcriptional level of the latter, a mechanism possibly relevant to functionally related genes that must be expressed at different strengths while keeping a constant ratio (59). The termination efficiency can also be subject to regulation, depending on environmental conditions and metabolic needs, resulting in a variable degree of read-through and thus of relative expression levels (57, 58). Such conditional read-through can involve Rho and other proteins assisting termination (60, 61, 62, 63) as well as other conditional premature termination mechanisms such as attenuation (64, 65), T-box conditional termination (66, 67) and riboswitches (68, 69).

An example of condition-independent read-through occurs at the *rhlB-gppA-pehV* TU (Fig. 4A and Supplementary Fig. S1C). The *rhlB* gene encodes a component of the RNA degradosome (70, 71) whereas *gppA* encodes guanosine 5’-triphosphate 3’-diphosphate (pppGpp) pyrophosphatase involved in bacterial stringent response (72) and the *pehV* gene encodes a polygalacturonase involved in pectin degradation (73). These genes are functionally unrelated (except for a distant link to nutritional stress) yet appear co-transcribed, which is in fact quite frequent among operons (41, 74). This TU exhibits a variable expression level (by up to 50%) across the sampled conditions, but the internal (relative) expression pattern is condition-independent: *rhlB* and *gppA* are expressed at a similar level, whereas *pehV* is systematically less transcribed (Fig. 4A and and Supplementary Fig. S1C). This observation is correlated with the presence of an intrinsic internal TTS downstream of *gppA.* By computing the expression ratio of *pehV* compared to *rhlBlgppA*, we inferred the associated termination probability (or terminator strength) and found a constant value *P*(*TTS_gppA_*) = 92 ± 1% (95% confidence interval) characteristic of a non-conditional transcriptional read-through. Thus, the three genes are co-transcribed from a single promoter of condition-dependent activity, with a reduced transcriptional level of *pehV* exhibiting a constant ratio (8%) compared to the other genes. The biological relevance of this mechanism remains to be clarified. In *E. coli, rhlB* and *gppA* were also recently shown to be co-transcribed (3, 54). Another example of condition-independent read-through occurs at the *gcvTHP* TU involved in glycine cleavage (75) (Fig. 6). We detected an internal TTS downstream of *gcvH* in accordance with studies in *E. coli* (3, 54) and inferred its termination probability *P*(*TTS_gcvH_*) = 71 ± 22% (95% confidence interval), based on the expression ratio of *gcvP* compared to *gcvT* and *gcvH* across RNA-seq conditions. It is unclear whether this variability is due to RNA-seq signal variations or a weak regulation of the termination rate. The GcvT, H, and P proteins are part of the glycine cleavage system with GcvL (76), and GcvP activity might be required at lower concentration in the investigated conditions.

By definition, all identified internal TTSs (549) experience transcriptional read-through. As a rough estimate, condition-independent read-through was detected for 77 (14%) of internal TTSs, based on the constant expression ratio of the genes located down-stream vs upstream across RNA-seq conditions (Fig. 4, Materials and Methods). The remaining internal TTSs rather experience condition-dependent read-through; however, the systematic estimation of stochastic termination rates at internal TTSs is delicate based on our data only, due to the limited number of RNA-seq conditions and the presence of nearby TSSs that contribute to the heterogeneous expression levels along the TU, as illustrated by *pelCZ* (Fig. 3D).

An example of condition-dependent read-through occurs at the *pelD-paeY-pemA* TU (Fig. 4B and Supplementary Fig. S2B), which is identified by our approach but was also characterised by Northern blotting (77). It encodes three genes involved in pectin degradation. In the initial step of pectinolysis occurring in plants, *paeY* (acetylesterase) and *pemA* (methylesterase) remove acetyl and methyl groups from pectin, which can then be efficiently degraded by the pectate lyase *pelD* (17). The *pelD* gene is essentially transcribed as a monocistronic RNA, although its terminator (predicted as intrinsic) can be overstepped to generate a polycistronic transcript comprising the three genes (77). In exponential phase, the three genes are homogeneously (but weakly) transcribed as a unique polycistronic RNA, suggesting that the internal TTS is not efficient (*P*(*TTS_pelD_*) = 0%). In stationary phase in presence of PGA, the whole TU is up-regulated, and the internal TTS becomes more efficient (*P*(*TTS_pelD_*) = 86 ± 3%, 95% confidence interval), resulting in the extensive synthesis of the *pelD* monocistronic RNA and a lower expression level of the two downstream genes. The regulation events occurring at this TTS remain to be characterised, but may adjust the relative expression levels of the genes following metabolic needs, since PelD has a predominant role in pectin degradation and virulence (78, 79) and must likely be required at much higher concentrations than the two other enzymes. In addition, the fact that *pemA* is differentially expressed depending on the degree of pectin methylation (80) highlights the relevance of adjusting the relative expression levels of the three genes depending on plant cell-wall composition.

Another example occurs at the *cytABCD* TU (Supplementary Fig. S2C and S3A). In addition to plants, *D. dadantii* is able to infect insects (81), during which this TU expresses four insecticidal toxins and was previously shown to produce a polycistronic mRNA comprising the four genes, besides the possible existence of alternative isoforms (82). The sequencing coverage together with the putative internal intrinsic TTS detected after *cytA* are clearly indicative of a condition-dependent read-through, with termination occurring less efficiently at *cytA* in stationary phase in presence of PGA compared to exponential phase. This variation in termination efficiency at *cytA* associated to an environmental change may again allow tuning the relative amounts of the corresponding toxins, especially if a precise and condition-dependent balance between them is required for optimal activity during the insect infection process (82). Interestingly, this cluster of four genes was acquired by horizontal transfer. Since transcriptional read-through partly relies on basal RNA Polymerase / TTS interactions, it might be conserved during horizontal transfer among bacterial species without requiring an independent acquisition of regulatory signals and their integration in the transcriptional regulatory network of the recipient cell.

### Detection of putative excludons and noncontiguous transcriptions units

All previous examples involved genes located on the same DNA strand; yet recent studies also describe interactions between overlapping antisense coding transcripts, involved in a mutual regulation. In particular, noncontiguous operons refer to operons that contain a gene or group of genes that is transcribed in the opposite direction (83). 83 TUs with such features were found in the *D. dadantii* genome (provided in Supplementary Tab. S4A). Among them, an example is the *indCvfmAZBCDFG* TU encoding a component of the *vfm* quorum sensing system required for the production of plant cell wall-degrading enzymes (Fig. 5 and Supplementary Fig. S2D) (84). The *vfmE* gene, located on the opposite strand and within this TU, is also part of this system and known to encode a transcriptional activator of the *vfm* locus (of the AraC family). Since all genes of the TU are co-transcribed within a single mRNA, it is likely that these two overlapping antisense transcripts could negatively regulate each other, *e.g*. by transcriptional interference (RNA Polymerases collision) or RnaseIII-mediated double-stranded RNA processing (85). An expression increase of the *vfm* locus would then reduce the expression of *vfmE*, and in turn its own expression, forming a genome-embedded negative feedback loop controlling the production of quorum sensing signal and plant cell-wall degrading enzymes (86).

**FIG 5.**
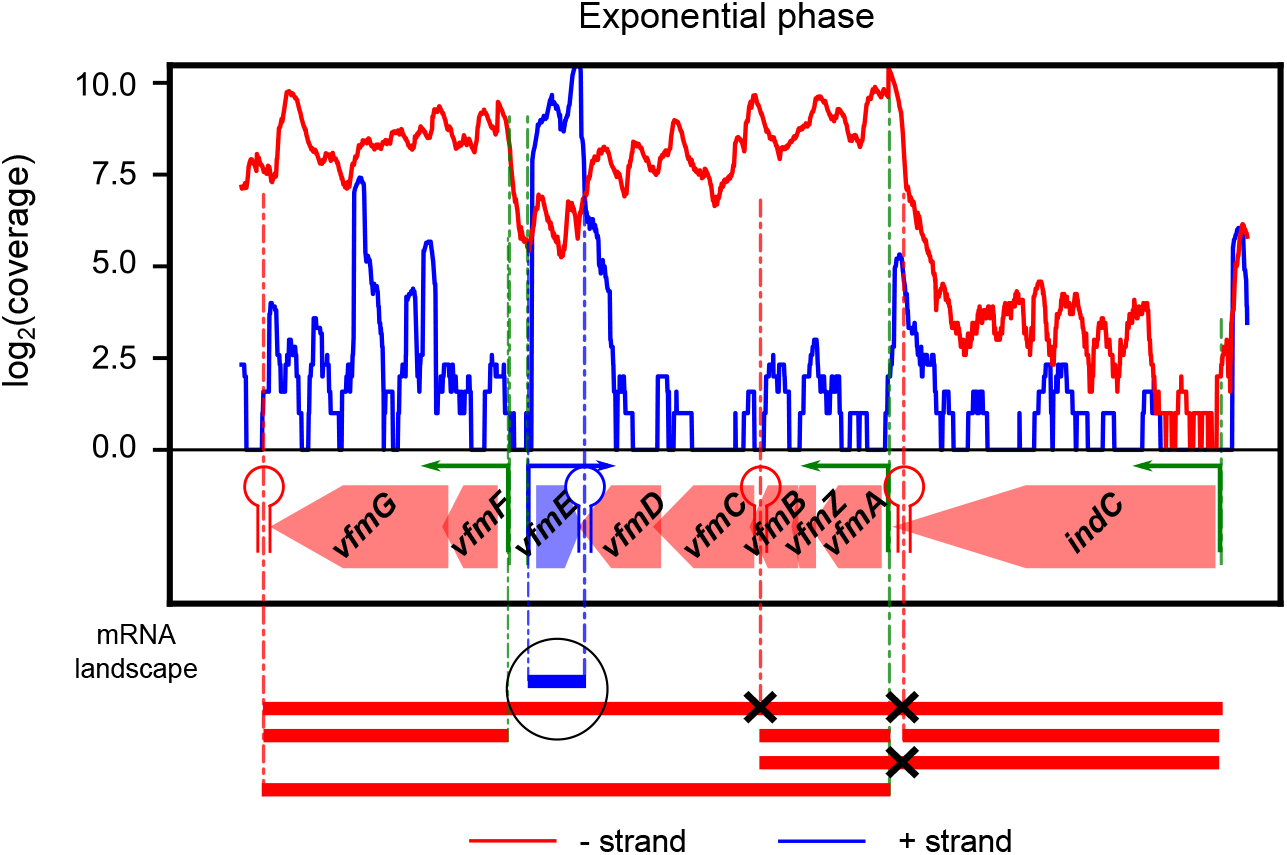
Existence of a potential noncontiguous transcription unit in the *vfm* locus. The *vfmE* gene is transcribed in the opposite direction of the *indCvfmAZBCDFG* TU, generating two overlapping mRNAs (as shown in red/blue for the -/+ strand) that might be involved in a mutual regulation (see text). All mRNA isoforms are observed in Nanopore native RNA-seq data (Supplementary Fig. S2D), including a long native RNA read on the negative strand between *vfmD* and *vfmF*.

**FIG 6.**
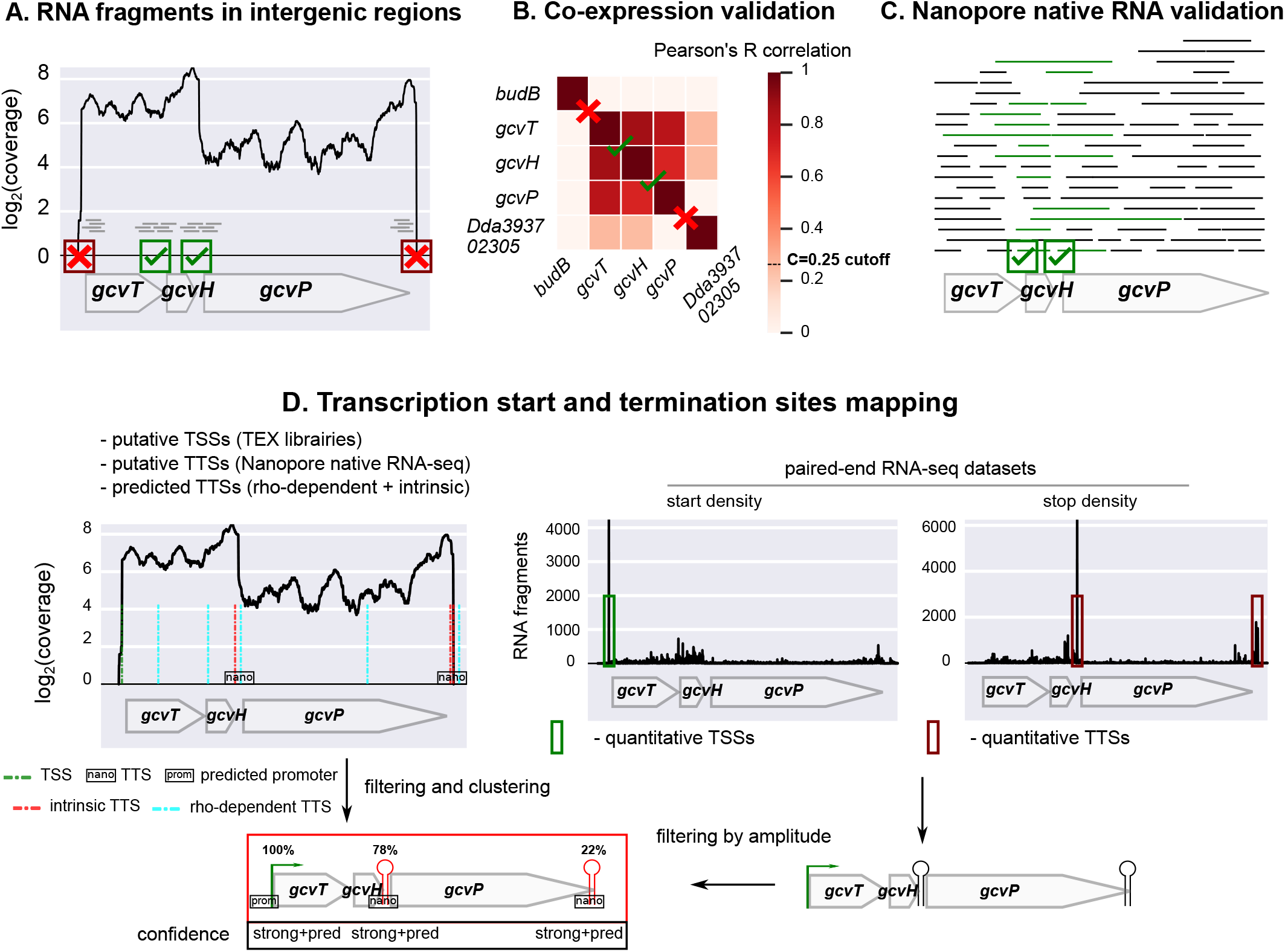
Algorithm for the characterisation of *D. dadantii* transcription units. (A) Definition of putative TUs based on RNA-seq coverage (dataset 1) in intergenic regions, with isodirectional genes being split when the coverage drops to zero (here before *gcvT* and after *gcvP*). (B) Validation based on correlation of expression across 32 conditions (dataset 2). The genes of identified putative TUs are correlated, in contrast to surrounding isodirectional genes (*budB* and *Dda3937_02305*). (C) Validation based on Nanopore native RNA-seq, based on the presence of overlapping RNA reads between adjacent gene pairs, yielding a direct evidence for co-transcription. (D) TSS and TTS mapping based on dRNA-seq (TEX libraries, dataset 4), Nanopore native RNA-seq (dataset 3), TTS predictions, promoter predictions, and paired-end RNA-seq data (dataset 1). First, putative TSSs and TTSs of high resolution but qualitative strength were defined from an analysis of TEX libraries and Nanopore native RNA-seq, respectively, and rho-dependent/intrinsic terminations were predicted. Second, a list of TSSs and TTSs of quantitative strength but poorer resolution was defined from the enrichment of RNA-seq paired-end fragment starts (start density) and stops (stop density) upstream of gene starts and downstream of gene stops, respectively. Third, only TSSs and TTSs with sufficient strength were retained, and compared to the closest TEX TSS / Nanopore TTS / predicted hairpin loop, in order to define their exact position and level of confidence. Finally, promoters were predicted for the retained TSSs. As a result of the analysis, this TU included the *gcvTHP* genes, the first two genes being expressed both as part of the entire transcript and as an independent transcript generated from a strong internal TTS (76% of total magnitude), explaining the lower correlation between *gcvP* and the remaining genes.

Finally, “excludons” refer to genomic regions in which convergent or divergent genes display overlapping transcription (87). From the map of transcription start and termination sites, we found 160 putative convergent excludons (overlapping 3’ UTRs) and 63 putative divergent excludons (overlapping 5’ UTRs) (provided in Supplementary Tab. S4B). An example is the divergent excludon between *greB* and *ompRenvZ* transcription units, encoding a transcript cleavage factor required for effective transcription elongation (88) and a two-component signal transduction system involved in osmotic stress response (89), respectively (Supplementary Fig. S3B). Both TUs comprise long 5’UTRs, forming a region of overlapping transcription that was previously identified in *E. coli* (90) and might underpin a mutual post-transcriptional regulation.

### *In planta* co-expression validation of the transcription units

While our transcriptional map was inferred from *in vitro* cultures, where RNAs could be extracted with optimal quality and reproducibility, we wished to test if the identified TUs could play a role in conditions of plant infection. We analysed a set of expression data obtained *in planta* by DNA microarrays, during the early stages of *Arabidopsis thaliana* infection (dataset 5) (29), 6 hours post-inoculation, during the epiphytic colonisation of leaf surface, and 24 hours post-inoculation during leaf invasion, just before the onset of visible symptoms. Overall, among the 50% gene pairs most correlated *in planta*, 80% belong to the same TUs, suggesting that co-transcription of these genes may indeed likely occur in these conditions (Supplementary Fig S4A). As an example, in *cytABCD*, the four genes are also highly correlated *in planta*, while this correlation immediately drops in surrounding isodirectional TUs (Supplementary Fig. S4B), as we expected. However, comparable correlations might also arise between other genes that are not transcribed together, but share the same transcriptional regulators, in particular those involved in virulence such as KdgR, PecT, PecS (91), thus accounting for the 20% strongly correlated gene pairs not located in the same TU. For example, in the *pelCZ* complex TU involved in pectinolysis, both genes are strongly correlated *in planta* (Supplementary Fig. S4C), as expected from the previous *in vitro* observations (especially with PGA, Supplementary Fig. S4C), but the adjacent *pelB* gene is also correlated, whereas *crp* and *mrcA* are not. This is not surprising, since most *pel* are paralogous genes with similar regulators and are strongly induced by pectin. The same pattern is observed for the *pelD-paeY-pemA* TU (Supplementary Fig. S4D), with respect to the *pelE* and *pelA* genes located upstream on the same strand. Because of the limited spatial resolution of microarrays and the weak number of investigated conditions, it is not possible to systematically distinguish the effects of these two mechanisms at the genomic scale from these data, but a survey of representative TUs confirmed that they usually coincide with correlated blocks of genes (as observed with *cytABCD*), even when the latter do not belong to the same functional pathways.

As an example, the complex TU *sufABCDSE-ldtC* is composed of two functionally unrelated operons (Supplementary Fig. S4E). *sufABCDSE* encodes components of the iron-sulfur cluster assembly machinery (92), which is required to synthesise and repair damaged iron-sulfur clusters under conditions of oxidative stress or iron limitation, and is therefore critical for *D. dadantii* virulence (91). In contrast, *ldtC* (previously *ycfS*), encodes a L,D-transpeptidase crucial for bacterial envelope assembly, by catalysing the attachment of the major outer-membrane protein Lpp to peptidoglycan (93). According to our findings above, *sufABCDSE* and *ldtC* can be transcribed together, with an internal TTS and TSS located between them. *In planta*, the seven genes are indeed strongly co-expressed, with a slight decrease for *ldtC*, in full agreement with the identified transcriptional map (Supplementary Fig S4E). It is conceivable that these genes are required under a common set of conditions encountered during plant infection, which was favoured by their inclusion in the same transcript, while the presence of alternative TSS and TTS might still allow separate expression when required. Indeed, the *sufABCDSE* operon is controlled by three transcriptional regulators, Fur, OxyR and IscR, which respectively sense iron limitation, oxidative stress and intracellular iron-sulfur cluster status (94). Each of them contributes to the activation of the *suf* promoter by oxidative stress occurring during plant penetration and colonisation (25): the repressor Fur is inactivated by reactive oxygen species (ROS); the activator OxyR becomes active through the oxidation of two cysteine residues and the formation of a disulfide bond; IscR becomes an activator of *suf* promoter after destruction of its iron-sulfur cluster by ROS (94). On the other hand, the activity of L,D-transpeptidases involves a catalytic cysteine residue that must be reduced (95), which is challenging under oxidative stress. The expression of *ldtC* from the *suf* promoter, which is strongly activated in the latter condition, is therefore biologically meaningful. Interestingly, in *E. coli*, the *suf* operon is also located upstream of a gene encoding a L,D-transpeptidase (*ldtA*), the two operons being also transcribed both together and separately (54).

### Concluding statement

In this study, we combined five transcriptomic datasets yielding complementary information and designed to provide a catalogue of genes’ responses and RNA landscapes to various growth and stress conditions, including one of the first applications of Nanopore native RNA-seq to prokaryotic transcriptomes. Their integration through a computational method developed for this study allowed us to precisely determine and annotate the transcriptomic map of *D. dadantii*, the first of its kind in the *Dickeya* genus. The analysis of *in planta* DNA microarray data suggests that the identified TUs are also co-expressed during the early stages of plant infection, although a more refined *in planta* analysis would require higher-resolution transcriptomic data. Beyond its practical aspect as a community resource to help the scientific community to unravel gene regulation, including the virulence programme of this and related species, the obtained transcriptional map clearly indicates, after others, that the canonical operon model is insufficient to account for the complexity of bacterial transcription. The ability of the cell to differentially express genes of the same operon depending on metabolic needs and environmental conditions was first described with suboperonic regulation years ago. Later, with the emergence of next-generation sequencing, transcriptomic analyses confirmed at the genomic scale that most operons were able to generate alternative transcripts of variable gene composition. Transcriptional read-through at terminators is another mechanism that might play a basal coordination and regulation role, and explain the extent of transcription beyond the scale of operons. Recent findings include noncontiguous operons and excludons, where the expression of an operon transcript can be mutually regulated with that of a gene located on the opposite strand at the same locus. For such features, the putative catalogue provided here may be used as a starting point for further investigation, and in particular, might be combined with the *D. dadantii* non-coding RNA landscape (96) for a comprehensive analysis of transcriptional regulation in this bacterium. Altogether, our findings provide insights into the mechanisms of basal coordination of transcription and might contribute to the revision of the canonical view of operon structure and transcription organisation.

## MATERIALS AND METHODS

### Bacterial strain, genome annotation and genome-wide predictions of operons

The genome sequence and annotation files from *Dickeya dadantii* strain 3937 were obtained from NCBI under accession NC_014500.1 (97). This work focused on coding genes only (CDS, representing 4211 genes over 4411 in total). *D. dadantii* operons were predicted using Rockhopper, a recent computational tool for operon prediction based on RNA-seq expression data as well as genomic and functional information (40), by providing dataset 1 as input.

### RNA-sequencing data (dataset 1), definition of putative transcription units based on intergenic signals, and identification of unannotated genes

Strand-specific, paired-end RNA-seq processed data used in this study are described in (98). Transcriptomes were obtained in 6 conditions (with two biological replicates each) including various growth (M63 medium supplemented with sucrose, in exponential or stationary phase, in presence or absence of PGA) and DNA supercoiling conditions (novobiocin shock). For each sample, RNA fragments were inferred from paired-end reads information, and genome-wide coverage was computed from resulting RNA fragments coordinates using a Python home-made script.

To define putative transcription units, separately for each strand, adjacent genes were fused in the same putative TU as long as the coverage was greater than 0 at each position of their intergenic region (independently of its size) for at least half of the samples (Fig. 6A).

Unannotated genes were defined as DNA regions outside of known coding sequences, longer than the first centile (1%) of *D. dadantii* gene lengths (192 bp), with an average coverage significantly different from 0 (with 99% confidence, *i.e.*, > 9 at each position) in all samples, and with a coding sequence predicted by Prodigal (99), resulting in 50 unannotated genes. A search for homolog proteins was performed using PSI-BLAST based on the non-redundant protein database (Supplementary Tab. S1D).

### *In vitro* DNA microarray data (dataset 2) and co-expression validation of the putative transcription units using hierarchical clustering

Microarray processed data used in this study are described elsewhere (27). They comprise 32 *in vitro* conditions (with two biological replicates each) including various growth and stress conditions encountered by *D. dadantii* during plant infection: cells were harvested in M63 (minimal) medium supplemented with sucrose, in exponential or stationary phase, in presence or in absence of PGA or leaf extract, and exposed or not to environmental perturbations (acidic, osmotic and oxidative stress). Pearson’s correlation coefficients were computed among all gene pairs over all conditions on the logarithm of the normalised expression level (derived from probes intensity). For each putative TU, adjacent genes were grouped into clusters based on this correlation, using a hierarchical clustering framework constrained to group adjacent genes only, with a custom Python script. At each iteration of the algorithm, the median of cross-correlations among all clusters (or genes) was computed, and the adjacent clusters with maximal median were fused. The hierarchical clustering ends when a cutoff value *C* for the correlation is reached (Fig. 6B). If the agglomeration of all genes of the TU is achieved without reaching *C*, the TU is validated. Otherwise, the final clusters are considered as separate TUs. A high *C* value results in short highly correlated TUs, whereas a low *C* value yields longer moderately correlated TUs (Supplementary Tab. S5). We defined the value *C* = 0.25 such that 20% of operon predictions were discarded (Supplementary Fig. S5), since it is the number of false predictions (*i.e*. specificity) evaluated for Rockhopper in *E. coli*, a *D. dadantii* enterobacterium relative. Varying the precise value of C did not qualitatively change the main results (Supplementary Tab. S5). The identified TUs exhibit a similar length distribution as those reported in *E. coli* (9, 3).

### Nanopore native RNA sequencing (dataset 3), validation of the mRNA landscape, and genome-wide identification of putative transcription termination sites

*D. dadantii* cultures were grown in M63 medium supplemented with 0.2% glucose and 0.2% PGA, until the early exponential phase (*A*_600*nm*_ = 0.2, condition 1), or the early stationary phase (*A*_600*nm*_ = 1.8, condition 2). RNAs were extracted using a frozen acid-phenol method, as previously described (100), and treated successively with Roche and Biolabs DNases. Two samples were prepared: 50 μg of RNAs from each condition were pulled into one sample (sample 1), whereas the other one contained 100 μg of RNAs from condition 2 (sample 2). Both samples were then supplied to Vertis Biotechnologie AG for Nanopore native RNA-seq: total RNA preparations were first examined by capillary electrophoresis, and ribosomal RNA molecules were depleted for sample 1 only using an in-house developed protocol (recovery rate = 84%). RNA 3’ends were then poly(A)-tailed using poly(A) polymerase, and the Direct RNA sequencing kit (SQK-RNA002) was used to prepare the library for 1D sequencing on the Oxford Nanopore sequencing device. The direct RNA libraries were sequenced on a MinION device (MIN-101B) using standard settings. Basecalling of the fast5 files was performed using Guppy (version 3.6.1) with the following settings: -flowcell FLO-MIN106 - kit SQK-RNA002 -cpu_threads_per_caller 12-compress_fastq-reverse_sequence true-trim_strategy rna. Reads smaller than 50 nt were removed. 466 393 and 556 850 reads were generated from sample 1 and 2, respectively. Raw read sequencing data are available in the EBI Gene Expression (ArrayExpress) database under accession E-MTAB-10482. Quality control was performed on both datasets using Nanopack (101). Long-reads from the fastq files were mapped to *Dickeya dadantii* strain 3937 genome (NCBI accession number: NC_014500.1) (97) using minimap2 (release minimap2-2.17 (r941)) (102). Output alignments in PAF and SAM format were generated with the recommended options for noisy Nanopore native RNA-seq, adapted to bacteria (no splicing) (-ax map-ont-k14). Secondary alignments were not reported for sample 2 due to multiple secondary alignments in ribosomal RNAs regions (-secondary=no). In total, 382 290 and 392 743 alignments were generated (77 and 67% mappability) from sample 1 and 2, respectively. Alignment files were further sorted, indexed and analysed with SAMtools. Alignments from both samples were merged into one PAF file, and the latter was used for further analyses.

For each TU previously defined with datasets 1 and 2 (Fig. 6A and 6B), the presence of long overlapping native RNA reads was investigated using a Python home-made script for adjacent gene pairs belonging to the same TU (Fig. 6C). If at least one RNA read overlapped the two adjacent genes, their co-transcription was validated (quoted “validated” in Supplementary Tab. S1A). If the signal was too weak for the investigated genes (read counts <9, not significantly different from 0 with 99% confidence), no conclusion could be drawn (quoted “weak signal” in Supplementary Tab. S1A). Otherwise, if no overlapping RNA was found, it was not validated (quoted “invalidated” in Supplementary Tab. S1A), which might also be due to the low number of conditions tested.

For the determination of TTSs, for each position of the genome, we computed the total number of RNA fragments ending at this particular position, using a Python home-made script. From this stop density, we defined putative TTSs as positions downstream of gene stop codons (up to 100 bp, based on 3’UTR lengths in *E. coli*) enriched for RNA fragments stops, respectively. In each of these regions, we started from site *i* with highest stop signal *k_i_* on 5-bp centred windows (due to the low sequencing depth). For the position *i* to be considered as a putative TTS, we imposed *k_i_* to be significantly different from 0 (with 95% confidence, > 6). TTSs obtained with this approach are provided in Supplementary Tab. S2D.

### Differential RNA-sequencing experiments and genome-wide identification of putative transcription start sites (dataset 4)

RNAs from dataset 2 (27) (*in vitro* DNA microarray data) were pooled into four samples S1 to S4, resulting in a combination of stress (*pH*, *NaCl*, *H*_2_*O*_2_) and growth conditions: exponential phase with (S1) or without (S2) stress, transition to stationary phase with (S3) or without (S4) stress. Those samples were then supplied to Vertis Biotechnologie AG for TEX treatment and Illumina sequencing. Briefly, ribosomal RNA molecules were depleted from the total RNA samples using the Ribo-Zero rRNA Removal Kit for bacteria (Epicentre), and small RNAs (< 200 nt) were discarded using the RNeasy MinElute Cleanup Kit (Qiagen). For the generation of TSS cDNA libraries, the samples were first fragmented using RNase III, poly(A)-tailed using poly(A) polymerase, split into two halves, with one half being treated with Terminator exonuclease (+TEX, Epicentre), while the other one was left untreated (-TEX). 5’PPP structures were then converted into 5’P ends using RNA 5’ Polyphosphatase (5’PP, Epicentre), to which RNA adapters were ligated. First-strand cDNAs were synthetised using an oligo(dT)-adapter primer and the M-MLV reverse transcriptase, PCR-amplified using a high fidelity DNA polymerase, purified using the Agencourt AMPureXP kit (Beckman Coulter Genomics), and sequenced on an Illumina NextSeq 500 system (60 bp read length, single-end, strand-specific protocol). Sequencing reads were trimmed to remove poly(A) tails and adapters. The fastq sequencing files are available in the EBI Gene Expression (ArrayExpress) database under accession E-MTAB-9075. Putative TSS positions were then determined based on the enrichment of sequencing reads in TEX-treated samples (+TEX) compared to non-treated ones (-TEX) using TSSer, an automated annotation programme from dRNA-seq data with default parameters: TSS positions within 5 bases on the same strand were clustered together and the position with the highest amount of read increase in the +TEX library was retained. TSSs obtained with such approach are provided in Supplementary Tab. S2A.

### *In planta* DNA microarray data (dataset 5) and co-expression validation of the transcription units inferred from *in vitro* conditions

Microarray processed data used in this study are described in (29). They comprise two conditions: bacteria were collected 6 hours post-inoculation of the model plant *Arabidopsis thaliana* by wild-type *D. dadantii* during the epiphytic colonisation of the leaf surfaces (5 replicates), and 24 hours post-inoculation during the leaf invasion (4 replicates). Pearson’s correlation coefficient was computed among all gene pairs over the two conditions on the logarithm of the normalised expression level (derived from probes intensity) (Supplementary Fig. S4).

### Genome-wide detection of transcription start and termination sites from RNA-seq data, mapping to the transcription units

We computed the densities of RNA fragments starting and ending at each position of the genome, across all RNA-seq samples (dataset 1) (Fig. 6D). In order to retain only TSSs and TTSs relevant to proteincoding genes, we focused on regions located upstream of gene start codons (up to 250 bp, based on 5’UTR lengths in *E. coli*), and downstream of gene stop codons (up to 100 bp, based on 3’UTR lengths in *E. coli*), respectively. In each of these regions, putative TSSs/TTSs were defined as sites *i* with highest start/stop signal *k_i_*. To differentiate a TSS/TTS at position *i* from the noise, we imposed two successive conditions: (i) *k_i_* is significantly different from 0 (with 99% confidence, *k_i_* > 9); (ii) *k_i_* is greater or equal than a density cutoff value *D*. The latter was set as ten times the median of the density values of the region investigated for TSSs, and five times for TTSs, showing that the recorded transcripts indeed start/stop at that precise position, rather than along a poorly defined starting/stopping region. In that case, the position *i* was considered as a putative TSS/TTS, of strength *k_i_*. Setting a low density cutoff D would tend to include false positives resulting from RNA-seq signal variations (noise), whereas a high cutoff would exclude weakly expressed TSSs/TTSs. We selected the value of *D* (i) such that TSSs and TTSs were detected for known operons and experimentally characterised TUs (described in the manuscript) and (ii) by visually curating the density graphs and excluding many positions obviously associated to RNA-Seq signal variations.

TSSs/TTSs positions were then compared among datasets to evaluate their confidence level. For each TSS identified with this approach, if a putative TSS obtained from dataset 4 (TEX libraries) was close enough (±20*bp*), its position was retained (assuming a higher precision and resolution). In addition, a scan for promoter motifs was conducted with bTSSfinder (52). For TTSs, the same method was applied using the position of the closest predicted hairpin loop (±50 bp), or TTS positions obtained from Nanopore native RNA-seq data (dataset 3). TSSs and TSSs were then assigned to the TUs, and only internal TSSs and TTSs with 15% relative amplitude (*i.e.* 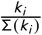) were retained, resulting in a total of 2595 TSSs and 1699 TTSs over all TUs. Setting a low relative amplitude cutoff would tend to retain all TSSs / TTSs, including many very weak ones mostly due to noise. We selected the relative amplitude cutoff value (i) based on a collection of known operons and TUs (shown in the manuscript), and (ii) such that the total number of TSSs and TTSs identified was consistent with those reported recently in *E. coli* (3, 54). If no TSS/TTS was found from dataset 1, we indicated the closest putative one from dataset 3/4 with a lower confidence level. The lists are provided in Supplementary Tab. S1A to S1C.

### Detection of transcriptional readthrough at internal TTSs

For each internal TTS, the expression ratio Δ(*log*_2_ (*RPKM*)) of the gene located downstream compared to the gene located upstream was computed across RNA-seq conditions. We imposed two successive conditions to consider the transcriptional read-through at this TTS as condition-independent: (1) Δ(*log*_2_(*RPKM*)) ≤ −0.5 for at least 8 samples over 12 corresponding at least to a termination probability *P*(*TTS*) = 71%, (2) standard error of the mean *σ P*(*TTS*) ≤ 12.5% corresponding to a relatively constant mean expression ratio and subsequent termination probability.

## Availability of data and materials

- *Dickeya dadantii* strain 3937 genome sequence and annotation files: NCBI accession number NC_014500.1 (97).
- RNA-seq data (dataset 1): EBI Gene Expression (ArrayExpress) accession number E-MTAB-7650 (98).
- *In vitro* microarray data (dataset 2): EBI Gene Expression (ArrayExpress) accession number E-MTAB-541 (27).
- Nanopore native RNA sequencing (dataset 3): EBI Gene Expression (ArrayExpress) accession number E-MTAB-10482. Note: not publicly available until the manuscript is accepted. Please use login details Username = Reviewer_E-MTAB-10482, Password =gdrof3hg.
- Differential RNA-seq data (dataset 4): EBI Gene Expression (ArrayExpress) accession number E-MTAB-9075. Note: not publicly available until the manuscript is accepted. Please use login details Username = Reviewer_E-MTAB-9075, Password = jenuxmon.
- *In planta* microarray data (dataset 5): NCBI Gene Expression Omnibus (GEO) accession number GSE94713 (29).

## SUPPLEMENTAL MATERIAL

**SUPPLEMENTARY FIGURE S1.** (A) Co-expression validation of *znuCBA* TU with *in vitro* DNA microarray data (dataset 2): the three genes exhibit strong internal cross-correlations clearly indicative of an operon. (B) Same for *sapABCDFfabI* TU: the six genes are coexpressed, with a reduced correlation of *fabI* due to the presence of a strong internal TSS (Fig. 3B). (C) Same for *rhlb-gppA-pehV* TU: the three genes are co-expressed, with a reduced transcriptional level of *pehV* (Fig. 4A) and a reduced correlation due to condition-independent read-through at the *gppA* intrinsic terminator. (D) Effect of PGA on *pelCZ* TU: *pelC* and *pelZ* expression profiles are similar in absence (left) or presence (right) of PGA in stationary phase, in spite of a drastically different global expression level.

**SUPPLEMENTARY FIGURE S2.** Co-transcription and mRNA landscape validation with Nanopore native RNA-seq for (A) *rhlB-gppA-pehV* TU with condition-independent read-through at the intrinsic TTS downstream of *gppA;* (B) *pelD-paeY-pemA* TU with condition-dependent read-through downstream of *pelD* internal TTS; (C) *cytABCD* TU with condition-dependent read-through downstream of *cytA* internal TTS. Those internal TTSs are occasionally overstepped, resulting in different transcripts isoforms (as shown in red) which are all detected as long native RNA reads (black). (D) *indCvf-mAZBCDFG* noncontiguous TU (black Nanopore reads on the negative strand), with *vfmE* being transcribed on the opposite strand (blue Nanopore reads on the positive strand), resulting in overlapping antisense transcripts.

**SUPPLEMENTARY FIGURE S3.** (A) Quantification of condition-dependent transcriptional read-through: example of the *cytABCD* TU. A putative Rho-independent TTS is identified downstream of *cytA* although not validated. Its probability of termination (inferred from the expression variation Δ(*log*_2_(*RPKM*)) of *cytA* compared to the other genes) is regulated and depends both on the growth phase and the presence of PGA (*P*(*TTS_cytA_*) = 0.78 ± 0.03 vs 0.51 ± 0.03) besides a global up-regulation of the whole TU. (B) The *greB* and *ompRenvZ* transcription units form a potential divergent excludon: long 5’UTRs overlapping transcripts are generated by *ompR* and *greB* divergent genes and might form a dsRNA that could prevent each other transcription. In *E. coli, ompR* and *envZ* are part of the same operon (red), and *greB* is transcribed alone (blue). Such genomic region forming a dsRNA was also identified in *E. coli* (90).

**SUPPLEMENTARY FIGURE S4.** *In planta* DNA microarray data, 6 hours post-inoculation (hpi) of the plant *Arabidopsis thaliana* (epiphytic colonisation of the leaf surfaces, 5 replicates), and 24 hpi (leaf invasion, 4 replicates). (A) Distribution of co-expression correlation coefficients among (blue) all genes and (red) genes belonging to the same TU. Among the 50% most correlated genes *in planta*, 80% belong to the same TUs, with the example TUs from the manuscript (*smtA-mukFEB, znuCBA, sapABCDF-fabI,glgBXCAP, pelCZ, rhlB-gppA-pehV, pelD-paeY-pemA, cytABCD)* displaying a median correlation of 0.9 *in planta*. (B) Pearson’s co-expression correlation coefficients of *cytABCD* TU with surrounding isodirectional (on the same strand) TUs. (C) Same for *pelCZ* TU. (D) Same for *pelD-paeY-pemA* TU. (E) Identification of *sufABCDSE-ldtC* complex TU, composed of operons of apparently unrelated functions, exhibiting a strong internal TSS (51% total magnitude) upstream of *ldtC* (previously *ycfS*) and a strong internal TTS (52% total magnitude) downstream *sufE*, allowing separate transcriptions. The seven genes are highly correlated *in planta*.

**SUPPLEMENTARY FIGURE S5.** Co-expression validation of transcription units for different correlation thresholds *C*. TUs obtained with high *C* values are more highly correlated but shorter. Putative TUs are obtained from step 1 of the analysis (intergenic signal), without any requirement on the correlation of expression. With the chosen value (*C* = 0.25), TUs group around three times more gene pairs than predicted operons by Rockhopper. The value of *C* was chosen such that 20% of operon predictions were discarded, since it is the number of false predictions of Rockhopper in *E. coli,* a *D. dadantii* enterobacterium relative.

**SUPPLEMENTARY TABLE S1.** (A) *Dickeya dadantii* transcription units defined by our approach. (B) TSSs across TUs. (C) TTSs across TUs. (D) Unannotated protein-coding genes.

**SUPPLEMENTARY TABLE S2.** (A) Putative TSSs identified by differential RNA-seq (TEX treatment) under a wide range of environmental conditions. (B) Genomic position and secondary structure of putative TTSs: intrinsic terminators predicted by ARNold (Erpin and RNAmotif algorithms). (C) Genomic position of putative TTSs: Rho-dependent terminators predicted by RhoTermPredict. (D) Putative TTSs identified by Nanopore native RNA-seq.

**SUPPLEMENTARY TABLE S3.** TSS validation, based on all published TSSs to date and to our knowledge in *D.dadantii.*

**SUPPLEMENTARY TABLE S4.** (A) Catalogue of putative noncontiguous transcription units. (B) Catalogue of putative excludons.

**SUPPLEMENTARY TABLE S5.** Catalogue of transcription unit architecture. Putative TUs are obtained from the first step of the approach (analysis of intergenic signal). Varying the precise value of the correlation threshold *C* for co-expression validation (step 2) does not change the results qualitatively. A larger *C* value results in shorter but more highly correlated TUs. Final TUs obtained with *C* = 0.25 are longer than predicted operons and exhibit a similar length distribution as those reported in *E. coli* (9, 3).

## ACKNOWLEDGEMENTS

We thank the whole CRP team for useful discussions as well as Ivan Junier.

## REFERENCES

1. Jacob F, Monod J. 1961. Genetic regulatory mechanisms in the synthesis of proteins. J Mol Biol 3:318–356. doi:10.1016/s0022-2836(61)80072-7.

2. Nicolas P, Mäder U, Dervyn E, Rochat T, Leduc A, Pigeonneau N, Bidnenko E, Marchadier E, Hoebeke M, Aymerich S, Becher D, Bisicchia P, Botella E, Delumeau O, Doherty G, Denham EL, Fogg MJ, Fromion V, Goelzer A, Hansen A, Härtig E, Harwood CR, Homuth G, Jarmer H, Jules M, Klipp E, Le Chat L, Lecointe F, Lewis P, Liebermeister W, March A, Mars RAT, Nannapaneni P, Noone D, Pohl S, Rinn B, Rügheimer F, Sappa PK, Samson F, Schaffer M, Schwikowski B, Steil L, Stülke J, Wiegert T, Devine KM, Wilkinson AJ, van Dijl JM, Hecker M, Völker U, Bessières P, Noirot P. Mar 2012. Condition-dependent transcriptome reveals high-level regulatory architecture in Bacillus subtilis. Sci (New York, N.Y.) 335 (6072):1103–1106. doi:10.1126/science.1206848.

3. Yan B, Boitano M, Clark TA, Ettwiller L. Sep 2018. SMRT-Cappable-seq reveals complex operon variants in bacteria. Nat Commun 9(1):1–11. doi:10.1038/s41467-018-05997-6.

4. Warrier I, Ram-Mohan N, Zhu Z, Hazery A, Echlin H, Rosch J, Meyer MM, Opijnen Tv. Dec 2018. The Transcriptional landscape of Streptococcus pneumoniae TIGR4 reveals a complex operon architecture and abundant riboregulation critical for growth and virulence. PLOS Pathog 14 (12):e1007461. doi:10.1371/journal.ppat.1007461.

5. Mejía-Almonte C, Busby SJW, Wade JT, van Helden J, Arkin AP, Stormo GD, Eilbeck K, Palsson BO, Galagan JE, Collado-Vides J. Jul 2020. Redefining fundamental concepts of transcription initiation in bacteria. Nat Rev Genet p 1–16. doi:10.1038/s41576-020-0254-8.

6. Adhya S. Jun 2003. Suboperonic regulatory signals. Sci STKE: signal transduction knowledge environment 2003 (185):pe22. doi: 10.1126/stke.2003.185.pe22.

7. Kornblihtt AR, Schor IE, Alló M, Dujardin G, Petrillo E, Muñoz MJ. Mar 2013. Alternative splicing: a pivotal step between eukaryotic transcription and translation. Nat Rev Mol Cell Biol 14 (3):153–165. doi: 10.1038/nrm3525.

8. Junier I, Rivoire O. 2016. Conserved Units of Co-Expression in Bacterial Genomes: An Evolutionary Insight into Transcriptional Regulation. PLOS One 11 (5):e0155740. doi:10.1371/journal.pone.0155740.

9. Conway T, Creecy JP, Maddox SM, Grissom JE, Conkle TL, Shadid TM, Teramoto J, San Miguel P, Shimada T, Ishihama A, Mori H, Wanner BL. 2014. Unprecedented high-resolution view of bacterial operon architecture revealed by RNA sequencing. mBio 5 (4):01442–01414. doi:10.1128/mBio.01442-14.

10. Kröger C, Dillon SC, Cameron ADS, Papenfort K, Sivasankaran SK, Hokamp K, Chao Y, Sittka A, Hébrard M, Händler K, Colgan A, Leekitcharoenphon P, Langridge GC, Lohan AJ, Loftus B, Lucchini S, Ussery DW, Dorman CJ, Thomson NR, Vogel J, Hinton JCD. May 2012. The transcriptional landscape and small RNAs of Salmonella enterica serovar Typhimurium. Proc Natl Acad Sci United States Am 109 (20):E1277–1286. doi:10.1073/pnas.1201061109.

11. Dugar G, Herbig A, Förstner KU, Heidrich N, Reinhardt R, Nieselt K, Sharma CM. May 2013. High-resolution transcriptome maps reveal strain-specific regulatory features of multiple Campylobacter jejuni isolates. PLOS genetics 9 (5):e1003495. doi: 10.1371/journal.pgen.1003495.

12. Wang Y, Li X, Mao Y, Blaschek HP. Sep 2011. Single-nucleotide resolution analysis of the transcriptome structure of Clostridium beijerinckii NCIMB 8052 using RNA-Seq. BMC Genom 12:479. doi: 10.1186/1471-2164-12-479.

13. Uplekar S, Rougemont J, Cole ST, Sala C. Jan 2013. High-resolution transcriptome and genome-wide dynamics of RNA polymerase and NusA in Mycobacterium tuberculosis. Nucleic Acids Res 41 (2):961–977. doi:10.1093/nar/gks1260.

14. Güell M, van Noort V, Yus E, Chen WH, Leigh-Bell J, Michalodimitrakis K, Yamada T, Arumugam M, Doerks T, Kühner S, Rode M, Suyama M, Schmidt S, Gavin AC, Bork P, Serrano L. Nov 2009. Transcriptome complexity in a genome-reduced bacterium. Sci (New York, N.Y.) 326 (5957):1268–1271. doi:10.1126/science.1176951.

15. Schmidtke C, Findeiss S, Sharma CM, Kuhfuss J, Hoffmann S, Vogel J, Stadler PF, Bonas U. Mar 2012. Genome-wide transcriptome analysis of the plant pathogen Xanthomonas identifies sRNAs with putative virulence functions. Nucleic Acids Res 40 (5):2020–2031. doi: 10.1093/nar/gkr904.

16. Alkhateeb RS, Vorhölter FJ, Rückert C, Mentz A, Wibberg D, Hublik G, Niehaus K, Pühler A. May 2016. Genome wide transcription start sites analysis of Xanthomonas campestris pv. campestris B100 with insights into the gum gene cluster directing the biosynthesis of the exopolysaccharide xanthan. J Biotechnol 225:18–28. doi: 10.1016/j.jbiotec.2016.03.020.

17. Hugouvieux-Cotte-Pattat N, Condemine G, Gueguen E, Shevchik VE. 2020. Dickeya Plant Pathogens, p 1–10. In eLS. American Cancer Society. doi:10.1002/9780470015902.a0028932.

18. Fujikawa T, Ota N, Sasaki M, Nakamura T, Iwanami T. Jul 2019. Emergence of apple bacterial quick decline caused by Dickeya dadantii in Japan. J Gen Plant Pathol 85 (4):314–319. doi:10.1007/s10327-019-00852-y.

19. Toth IK, Wolf JMvd, Saddler G, Lojkowska E, Hélias V, Pirhonen M, Tsror (Lahkim) L, Elphinstone JG. 2011. Dickeya species: an emerging problem for potato production in Europe. Plant Pathol 60 (3):385–399. doi:10.1111/j.1365-3059.2011.02427.x.

20. Jiang HH, Hao JJ, Johnson SB, Brueggeman RS, Secor G. Jul 2016. First Report of Dickeya dianthicola Causing Blackleg and Bacterial Soft Rot on Potato in Maine. Plant Dis 100 (11):2320–2320. doi: 10.1094/PDIS-12-15-1513-PDN.

21. Pu XM, Zhou JN, Lin BR, Shen HF. Dec 2012. First Report of Bacterial Foot Rot of Rice Caused by a Dickeya zeae in China. Plant Dis 96 (12):1818. doi:10.1094/PDIS-03-12-0315-PDN.

22. Ma B, Hibbing ME, Kim HS, Reedy RM, Yedidia I, Breuer J, Breuer J, Glasner JD, Perna NT, Kelman A, Charkowski AO. Sep 2007. Host range and molecular phylogenies of the soft rot enterobacterial genera pectobacterium and dickeya. Phytopathology 97 (9):1150–1163. doi:10.1094/PHYTO-97-9-1150.

23. Reverchon S, Muskhelisvili G, Nasser W. 2016. Virulence Program of a Bacterial Plant Pathogen: The Dickeya Model. Prog Mol Biol Transl Sci 142:51–92. doi:10.1016/bs.pmbts.2016.05.005.

24. Grignon C, Sentenac H. 1991. pH and Ionic Conditions in the Apoplast. Annu Rev Plant Physiol Plant Mol Biol 42 (1):103–128. doi: 10.1146/annurev.pp.42.060191.000535.

25. Lamb C, Dixon RA. 1997. The Oxidative Burst in Plant Disease Resistance. Annu Rev Plant Physiol Plant Mol Biol 48 (1):251–275. doi: 10.1146/annurev.arplant.48.1.251.

26. Lebeau A, Reverchon S, Gaubert S, Kraepiel Y, Simond-Côte E, Nasser W, Van Gijsegem F. Mar 2008. The GacA global regulator is required for the appropriate expression of Erwinia chrysanthemi 3937 pathogenicity genes during plant infection. Environ Microbiol 10(3):545–559. doi:10.1111/j.1462-2920.2007.01473.x.

27. Jiang X, Sobetzko P, Nasser W, Reverchon S, Muskhelishvili G. Apr 2015. Chromosomal “Stress-Response” Domains Govern the Spatiotemporal Expression of the Bacterial Virulence Program. mBio 6 (3). doi:10.1128/mBio.00353-15.

28. Jiang X, Zghidi-Abouzid O, Oger-Desfeux C, Hommais F, Greliche N, Muskhelishvili G, Nasser W, Reverchon S. 2016. Global transcriptional response of Dickeya dadantii to environmental stimuli relevant to the plant infection. Environ Microbiol 18 (11):3651–3672. doi: 10.1111/1462-2920.13267.

29. Pédron J, Chapelle E, Alunni B, Van Gijsegem F. 2018. Transcriptome analysis of the Dickeya dadantii PecS regulon during the early stages of interaction with Arabidopsis thaliana. Mol Plant Pathol 19 (3):647–663. doi:10.1111/mpp.12549.

30. Duprey A, Taib N, Leonard S, Garin T, Flandrois JP, Nasser W, Brochier-Armanet C, Reverchon S. 2019. The phytopathogenic nature of Dickeya aquatica 174/2 and the dynamic early evolution of Dickeya pathogenicity. Environ Microbiol 21 (8):2809–2835. doi: 10.1111/1462-2920.14627.

31. Nasser W, Condemine G, Plantier R, Anker D, Robert-Baudouy J. Jun 1991. Inducing properties of analogs of 2-keto-3-deoxygluconate on the expression of pectinase genes of Erwinia chrysanthemi. FEMS Microbiol Lett 81 (1):73–78. doi:10.1111/j.1574-6968.1991.tb04715.x.

32. Ouafa ZA, Reverchon S, Lautier T, Muskhelishvili G, Nasser W. 2012. The nucleoid-associated proteins H-NS and FIS modulate the DNA supercoiling response of the pel genes, the major virulence factors in the plant pathogen bacterium Dickeya dadantii. Nucleic Acids Res 40 (10):4306–4319. doi:10.1093/nar/gks014.

33. Depledge DP, Srinivas KP, Sadaoka T, Bready D, Mori Y, Placantonakis DG, Mohr I, Wilson AC. Feb 2019. Direct RNA sequencing on nanopore arrays redefines the transcriptional complexity of a viral pathogen. Nat Commun 10 (1):754. doi:10.1038/s41467-019-08734-9.

34. Jenjaroenpun P, Wongsurawat T, Pereira R, Patumcharoenpol P, Ussery DW, Nielsen J, Nookaew I. Apr 2018. Complete genomic and transcriptional landscape analysis using third-generation sequencing: a case study of Saccharomyces cerevisiae CEN.PK113-7D. Nucleic Acids Res 46 (7):e38. doi:10.1093/nar/gky014.

35. Parker MT, Knop K, Sherwood AV, Schurch NJ, Mackinnon K, Gould PD, Hall AJ, Barton GJ, Simpson GG. Jan 2020. Nanopore direct RNA sequencing maps the complexity of Arabidopsis mRNA processing and m6A modification. eLife 9:e49658. doi: 10.7554/eLife.49658.

36. Workman RE, Tang AD, Tang PS, Jain M, Tyson JR, Razaghi R, Zuzarte PC, Gilpatrick T, Payne A, Quick J, Sadowski N, Holmes N, de Jesus JG, Jones KL, Soulette CM, Snutch TP, Loman N, Paten B, Loose M, Simpson JT, Olsen HE, Brooks AN, Akeson M, Timp W. Dec 2019. Nanopore native RNA sequencing of a human poly(A) transcriptome. Nat Methods 16(12):1297–1305. doi:10.1038/s41592-019-0617-2.

37. Pitt ME, Nguyen SH, Duarte TPS, Teng H, Blaskovich MAT, Cooper MA, Coin LJM. Feb 2020. Evaluating the genome and resistome of extensively drug-resistant Klebsiella pneumoniae using native DNA and RNA Nanopore sequencing. GigaScience 9 (giaa002). doi: 10.1093/gigascience/giaa002.

38. Sharma CM, Vogel J. Jun 2014. Differential RNA-seq: the approach behind and the biological insight gained. Curr Opin Microbiol 19:97–105. doi:10.1016/j.mib.2014.06.010.

39. Oyelade J, Isewon I, Oladipupo F, Aromolaran O, Uwoghiren E, Ameh F, Achas M, Adebiyi E. Nov 2016. Clustering Algorithms: Their Application to Gene Expression Data. Bioinform Biol Insights 10:237–253. doi:10.4137/BBI.S38316.

40. Tjaden B. Apr 2019. A computational system for identifying operons based on RNA-seq data. Methods doi:10.1016/j.ymeth.2019.03.026.

41. Yamanaka K, Ogura T, Niki H, Hiraga S. Nov 1995. Characterization of the smtA gene encoding an S-adenosylmethionine-dependent methyltransferase of Escherichia coli. FEMS microbiology letters 133 (1-2):59–63. doi:10.1111/j.1574-6968.1995.tb07861.x.

42. López-Solanilla E, García-Olmedo F, Rodríguez-Palenzuela P. Jun 1998. Inactivation of the sapA to sapF locus of Erwinia chrysanthemi reveals common features in plant and animal bacterial pathogenesis. The Plant Cell 10 (6):917–924. doi:10.1105/tpc.10.6.917.

43. Cronan JE, Thomas J. 2009. Bacterial Fatty Acid Synthesis and its Relationships with Polyketide Synthetic Pathways. Methods enzymology 459:395–433. doi:10.1016/S0076-6879(09)04617-5.

44. Bergler H, Fuchsbichler S, Högenauer G, Turnowsky F. Dec 1996. The enoyl-[acyl-carrier-protein] reductase (fabI) of Escherichia coli, which catalyzes a key regulatory step in fatty acid biosynthesis, accepts NADH and NADPH as cofactors and is inhibited by palmitoyl-CoA. Eur J Biochem 242 (3):689–694. doi:10.1111/j.1432-1033.1996.0689r.x.

45. Preiss J. Aug 2009. Glycogen: Biosynthesis and Regulation. EcoSal Plus 3 (2). doi:10.1128/ecosalplus.4.7.4.

46. Montero M, Almagro G, Eydallin G, Viale AM, Muñoz FJ, Bahaji A, Li J, Rahimpour M, Baroja-Fernández E, Pozueta-Romero J. Jan 2011. Escherichia coli glycogen genes are organized in a single glgBXCAP transcriptional unit possessing an alternative suboperonic promoter within glgC that directs glgAP expression. The Biochem J 433 (1):107–117. doi:10.1042/BJ20101186.

47. Patzer SI, Hantke K. Aug 2000. The zinc-responsive regulator Zur and its control of the znu gene cluster encoding the ZnuABC zinc uptake system in Escherichia coli. The J Biol Chem 275(32):24321–24332. doi: 10.1074/jbc.M001775200.

48. Jorjani H, Zavolan M. Apr 2014. TSSer: an automated method to identify transcription start sites in prokaryotic genomes from differential RNA sequencing data. Bioinform (Oxford, England) 30 (7):971–974. doi:10.1093/bioinformatics/btt752.

49. Ray-Soni A, Bellecourt MJ, Landick R. 2016. Mechanisms of Bacterial Transcription Termination: All Good Things Must End. Annu Rev Biochem 85 (1):319–347. doi:10.1146/annurev-biochem-060815-014844.

50. Naville M, Ghuillot-Gaudeffroy A, Marchais A, Gautheret D. Feb 2011. ARNold: a web tool for the prediction of Rho-independent transcription terminators. RNA biology 8 (1):11–13. doi: 10.4161/rna.8.1.13346.

51. Di Salvo M, Puccio S, Peano C, Lacour S, Alifano P. Mar 2019. RhoTermPredict: an algorithm for predicting Rho-dependent transcription terminators based on Escherichia coli, Bacillus subtilis and Salmonella enterica databases. BMC Bioinform 20 (1):117. doi: 10.1186/s12859-019-2704-x.

52. Shahmuradov IA, Mohamad Razali R, Bougouffa S, Radovanovic A, Bajic VB. Feb 2017. bTSSfinder: a novel tool for the prediction of promoters in cyanobacteria and Escherichia coli. Bioinformatics 33 (3):334–340. doi:10.1093/bioinformatics/btw629.

53. Silva SdAe, Echeverrigaray S. Nov 2012. Bacterial Promoter Features Description and Their Application on E. coli in silico Prediction and Recognition Approaches. Bioinformatics doi:10.5772/48149.

54. Ju X, Li D, Liu S. Nov 2019. Full-length RNA profiling reveals pervasive bidirectional transcription terminators in bacteria. Nat Microbiol 4 (11):1907–1918. doi:10.1038/s41564-019-0500-z.

55. Pissavin C, Robert-Baudouy J, Hugouvieux-Cotte-Pattat N. Dec 1996. Regulation of pelZ, a gene of the pelB-pelC cluster encoding a new pectate lyase of Erwinia chrysanthemi 3937. J Bacteriol 178 (24):7187–7196. doi:10.1128/jb.178.24.7187-7196.1996.

56. Garibaldi A, Bateman DF. Jan 1971. Pectic enzymes produced by Erwinia chrysanthemi and their effects on plant tissue. Physiol Plant Pathol 1 (1):25–40. doi:10.1016/0048-4059(71)90037-3.

57. Gusarov I, Nudler E. Nov 2001. Control of Intrinsic Transcription Termination by N and NusA: The Basic Mechanisms. Cell 107 (4):437–449. doi:10.1016/S0092-8674(01)00582-7.

58. Morita T, Ueda M, Kubo K, Aiba H. Aug 2015. Insights into transcription termination of Hfq-binding sRNAs of Escherichia coli and characterization of readthrough products. RNA 21 (8):1490–1501. doi: 10.1261/rna.051870.115.

59. Stringer AM, Currenti S, Bonocora RP, Baranowski C, Petrone BL, Palumbo MJ, Reilly AA, Zhang Z, Erill I, Wade JT. Feb 2014. Genome-ScaleAnalyses of Escherichia coli and Salmonella enterica AraC Reveal Noncanonical Targets and an Expanded Core Regulon. J Bacteriol 196 (3):660–671. doi:10.1128/JB.01007-13.

60. Boudvillain M, Figueroa-Bossi N, Bossi L. Apr 2013. Terminator still moving forward: expanding roles for Rho factor. Curr Opin Microbiol 16 (2):118–124. doi:10.1016/j.mib.2012.12.003.

61. Burns CM, Richardson LV, Richardson JP. May 1998. Combinatorial effects of NusA and NusG on transcription elongation and rho-dependent termination in Escherichia coli. J Mol Biol 278 (2):307–316. doi:10.1006/jmbi.1998.1691.

62. Mondal S, Yakhnin AV, Babitzke P. 2017. Modular Organization of the NusA- and NusG-Stimulated RNA Polymerase Pause Signal That Participates in the Bacillus subtilis trp Operon Attenuation Mechanism. J Bacteriol 199(14). doi:10.1128/JB.00223-17.

63. Lawson MR, Berger JM. Jun 2019. Tuning the sequence specificity of a transcription terminator. Curr Genet 65 (3):729–733. doi: 10.1007/s00294-019-00939-1.

64. Merino E, Yanofsky C. May 2005. Transcription attenuation: a highly conserved regulatory strategy used by bacteria. Trends genetics: TIG 21 (5):260–264. doi:10.1016/j.tig.2005.03.002.

65. Turnbough CL. 2019. Regulation of Bacterial Gene Expression by Transcription Attenuation. Microbiol molecular biology reviews: MMBR 83 (3). doi:10.1128/MMBR.00019-19.

66. Green NJ, Grundy FJ, Henkin TM. Jan 2010. The T box mechanism: tRNA as a regulatory molecule. FEBS letters 584 (2):318–324. doi: 10.1016/j.febslet.2009.11.056.

67. Zhang J, Chetnani B, Cormack ED, Alonso D, Liu W, Mondragón A, Fei J. Sep 2018. Specific structural elements of the T-box riboswitch drive the two-step binding of the tRNA ligand. eLife 7:e39518. doi: 10.7554/eLife.39518.

68. Millman A, Dar D, Shamir M, Sorek R. Jan 2017. Computational prediction of regulatory, premature transcription termination in bacteria. Nucleic Acids Res 45 (2):886–893. doi:10.1093/nar/gkw749.

69. Proshkin S, Mironov A, Nudler E. Oct 2014. Riboswitches in regulation of Rho-dependent transcription termination. Biochimica Et Bio-physActa 1839 (10):974–977. doi:10.1016/j.bbagrm.2014.04.002.

70. Coburn GA, Miao X, Briant DJ, Mackie GA. Oct 1999. Reconstitution of a minimal RNA degradosome demonstrates functional coordination between a 3’exonuclease and a DEAD-box RNA helicase. Genes & Dev 13 (19):2594–2603. doi:10.1101/gad.13.19.2594.

71. Py B, Higgins CF, Krisch HM, Carpousis AJ. May 1996. A DEAD-box RNA helicase in the Escherichia coli RNA degradosome. Nature 381 (6578):169–172. doi:10.1038/381169a0.

72. Hauryliuk V, Atkinson GC, Murakami KS, Tenson T, Gerdes K. May 2015. Recent functional insights into the role of (p)ppGpp in bacterial physiology. Nat Rev Microbiol 13 (5):298–309. doi: 10.1038/nrmicro3448.

73. Nasser W, Shevchik VE, Hugouvieux-Cotte-Pattat N. Nov 1999. Analysis of three clustered polygalacturonase genes in Erwinia chrysanthemi 3937 revealed an anti-repressor function for the PecS regulator. Mol Microbiol 34 (4):641–650. doi:10.1046/j.1365-2958.1999.01609.x.

74. de Lorenzo V, Danchin A. Sep 2008. Synthetic biology: discovering new worlds and new words. The new and not so new aspects of this emerging research field. EMBO Reports 9 (9):822–827. doi: 10.1038/embor.2008.159.

75. Stauffer LT, Fogarty SJ, Stauffer GV. May 1994. Characterization of the Escherichia coli gcv operon. Gene 142 (1):17–22. doi: 10.1016/0378-1119(94)90349-2.

76. Kikuchi G, Motokawa Y, Yoshida T, Hiraga K. Jul 2008. Glycine cleavage system: reaction mechanism, physiological significance, and hyperglycinemia. ProcJpn Acad Ser B, Phys Biol Sci 84 (7):246–263. doi: 10.2183/pjab/84.246.

77. Shevchik VE, Hugouvieux-Cotte-Pattat N. Jun 1997. Identification of a bacterial pectin acetyl esterase in Erwinia chrysanthemi 3937. Mol Microbiol 24 (6):1285–1301. doi:10.1046/j.1365-2958.1997.4331800.x.

78. Boccara M, Diolez A, Rouve M, Kotoujansky A. Jul 1988. The role of individual pectate lyases of Erwinia chrysanthemi strain 3937 in pathogenicity on saintpaulia plants. Physiol Mol Plant Pathol 33 (1):95–104. doi:10.1016/0885-5765(88)90046-X.

79. Beaulieu C. 1993. Pathogenic Behavior of Pectinase-Defective Erwinia chrysanthemi Mutants on Different Plants. Mol Plant-Microbe Interactions 6 (2):197. doi:10.1094/MPMI-6-197.

80. Dorel C, Hugouvieux-Cotte-Pattat N, Robert-Baudouy J, Lojkowska E. Jul 1996. Production ofErwinia chrysanthemi pectinases in potato tubers showing high or low level of resistance to soft-rot. Eur J Plant Pathol 102 (6):511–517. doi:10.1007/BF01877017.

81. Grenier AM, Duport G, Pagès S, Condemine G, Rahbé Y. Mar 2006. The Phytopathogen Dickeya dadantii (Erwinia chrysanthemi 3937) Is a Pathogen of the Pea Aphid. Appl Environ Microbiol 72 (3):1956–1965. doi:10.1128/AEM.72.3.1956-1965.2006.

82. Costechareyre D, Dridi B, Rahbé Y, Condemine G. Dec 2010. Cyt toxin expression reveals an inverse regulation of insect and plant virulence factors of Dickeya dadantii. Environ Microbiol 12 (12):3290–3301. doi:10.1111/j.1462-2920.2010.02305.x.

83. Sáenz-Lahoya S, Bitarte N, García B, Burgui S, Vergara-Irigaray M, Valle J, Solano C, Toledo-Arana A, Lasa I. 2019. Noncontiguous operon is a genetic organization for coordinating bacterial gene expression. Proc Natl Acad Sci United States Am 116 (5):1733–1738. doi: 10.1073/pnas.1812746116.

84. Nasser W, Dorel C, Wawrzyniak J, Van Gijsegem F, Groleau MC, Déziel E, Reverchon S. Mar 2013. Vfm a new quorum sensing system controls the virulence of Dickeya dadantii. Environ Microbiol 15 (3):865–880. doi:10.1111/1462-2920.12049.

85. Toledo-Arana A, Lasa I. 2020. Advances in bacterial transcriptome understanding: From overlapping transcription to the excludon concept. Mol Microbiol 113 (3):593–602. doi:10.1111/mmi.14456.

86. Baltenneck J, Reverchon S, Hommais F. Jan 2021. Quorum Sensing Regulation in Phytopathogenic Bacteria. Microorganisms 9 (2). doi: 10.3390/microorganisms9020239.

87. Sesto N, Wurtzel O, Archambaud C, Sorek R, Cossart P. Feb 2013. The excludon: a new concept in bacterial antisense RNA-mediated gene regulation. Nat Rev Microbiol 11 (2):75–82. doi: 10.1038/nrmicro2934.

88. Hsu LM, Vo NV, Chamberlin MJ. Dec 1995. Escherichia coli transcript cleavage factors GreA and GreB stimulate promoter escape and gene expression in vivo and in vitro. Proc Natl Acad Sci United States Am 92 (25):11588–11592. doi:10.1073/pnas.92.25.11588.

89. Cai SJ, Inouye M. Jul 2002. EnvZ-OmpR interaction and osmoregulation in Escherichia coli. The J Biol Chem 277 (27):24155–24161. doi: 10.1074/jbc.M110715200.

90. Lybecker M, Zimmermann B, Bilusic I, Tukhtubaeva N, Schroeder R. Feb 2014. The double-stranded transcriptome of Escherichia coli. Proc Natl Acad Sci United States Am 111 (8):3134–3139. doi: 10.1073/pnas.1315974111.

91. Reverchon S, Nasser W. Oct 2013. Dickeya ecology, environment sensing and regulation of virulence programme. Environ Microbiol Reports 5 (5):622–636. doi:10.1111/1758-2229.12073.

92. Nachin L, Loiseau L, Expert D, Barras F. Feb 2003. SufC: an unorthodox cytoplasmic ABC/ATPase required for [Fe-S] biogenesis under oxidative stress. The EMBO J 22 (3):427–437. doi: 10.1093/emboj/cdg061.

93. Magnet S, Bellais S, Dubost L, Fourgeaud M, Mainardi JL, Petit-Frère S, Marie A, Mengin-Lecreulx D, Arthur M, Gutmann L. May 2007. Identification of the l,d-Transpeptidases Responsible for Attachment of the Braun Lipoprotein to Escherichia coli Peptidoglycan. J Bacteriol 189 (10):3927–3931. doi:10.1128/JB.00084-07.

94. Roche B, Aussel L, Ezraty B, Mandin P, Py B, Barras F. Mar 2013. Iron/sulfur proteins biogenesis in prokaryotes: Formation, regulation and diversity. Biochimica et Biophys Acta (BBA) - Bioenerg 1827 (3):455–469. doi:10.1016/j.bbabio.2012.12.010.

95. Collet JF, Cho SH, Iorga BI, Goemans CV. Aug 2020. How the assembly and protection of the bacterial cell envelope depend on cysteine residues. The J Biol Chem 295 (34):11984–11994. doi: 10.1074/jbc.REV120.011201.

96. Leonard S, Meyer S, Lacour S, Nasser W, Hommais F, Reverchon S. Sep 2019. APERO: a genome-wide approach for identifying bacterial small RNAs from RNA-Seq data. Nucleic Acids Res 47 (15):e88. doi: 10.1093/nar/gkz485.

97. Glasner JD, Yang CH, Reverchon S, Hugouvieux-Cotte-Pattat N, Condemine G, Bohin JP, Van Gijsegem F, Yang S, Franza T, Expert D, Plunkett G, San Francisco MJ, Charkowski AO, Py B, Bell K, Rauscher L, Rodriguez-Palenzuela P, Toussaint A, Holeva MC, He SY, Douet V, Boccara M, Blanco C, Toth I, Anderson BD, Biehl BS, Mau B, Flynn SM, Barras F, Lindeberg M, Birch PRJ, Tsuyumu S, Shi X, Hibbing M, Yap MN, Carpentier M, Dassa E, Umehara M, Kim JF, Rusch M, Soni P, Mayhew GF, Fouts DE, Gill SR, Blattner FR, Keen NT, Perna NT. Apr 2011. Genome Sequence of the Plant-Pathogenic Bacterium Dickeya dadantii 3937. J Bacteriol 193 (8):2076–2077. doi: 10.1128/JB.01513-10.

98. El Houdaigui B, Forquet R, Hindré T, Schneider D, Nasser W, Reverchon S, Meyer S. 2019. Bacterial genome architecture shapes global transcriptional regulation by DNA supercoiling. Nucleic Acids Res 47 (11):5648–5657. doi:10.1093/nar/gkz300.

99. Hyatt D, Chen GL, Locascio PF, Land ML, Larimer FW, Hauser LJ. Mar 2010. Prodigal: prokaryotic gene recognition and translation initiation site identification. BMC Bioinform 11:119. doi:10.1186/1471-2105-11-119.

100. Hommais F, Oger-Desfeux C, Gijsegem FV, Castang S, Ligori S, Expert D, Nasser W, Reverchon S. Nov 2008. PecS Is a Global Regulator of the Symptomatic Phase in the Phytopathogenic Bacterium Erwinia chrysanthemi 3937. J Bacteriol 190 (22):7508–7522. doi: 10.1128/JB.00553-08.

101. De Coster W, D’Hert S, Schultz DT, Cruts M, Van Broeckhoven C. Aug 2018. NanoPack: visualizing and processing long-read sequencing data. Bioinformatics 34 (15):2666–2669. doi: 10.1093/bioinformatics/bty149.

102. Li H. Sep 2018. Minimap2: pairwise alignment for nucleotide sequences. Bioinformatics 34 (18):3094–3100. doi: 10.1093/bioinformatics/bty191.

