## Supplementary figures and images for "Mapping the complex transcriptional landscape of the phytopathogenic bacterium *Dickeya dadantii*"

### Supplementary Fig. S1

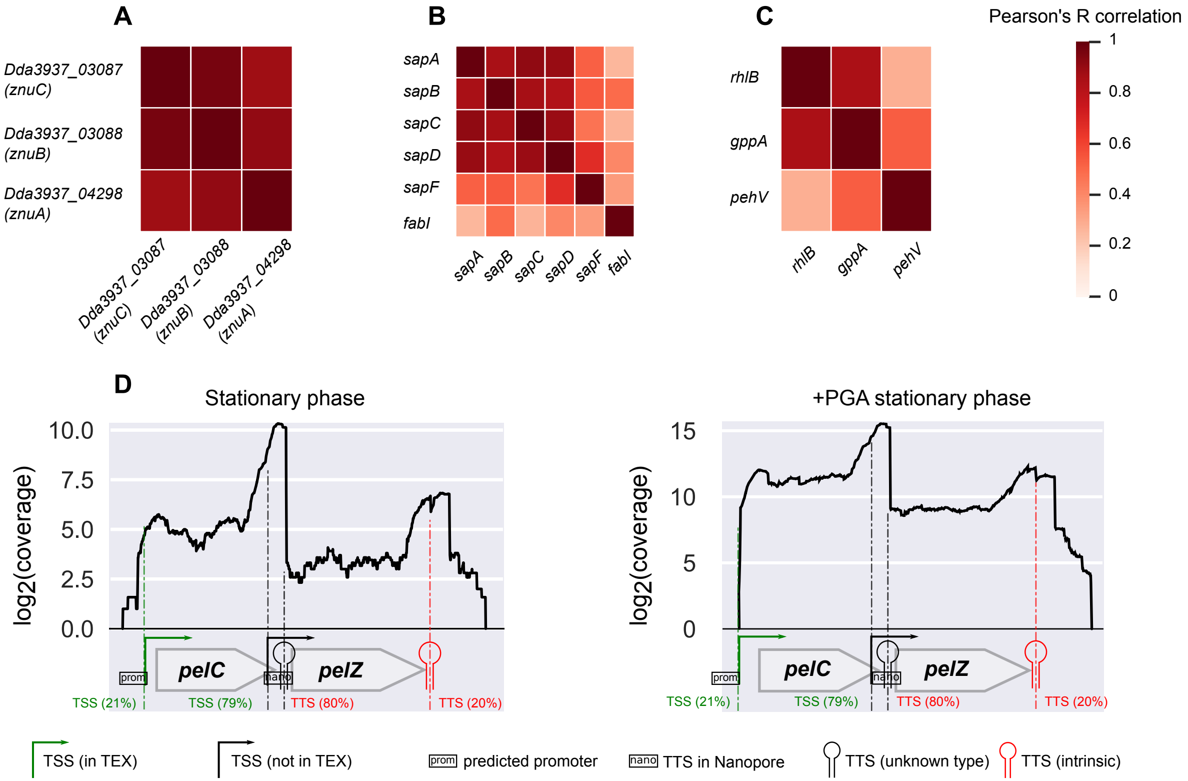

### Supplementary Fig. S2

#### Nanopore native RNA-seq

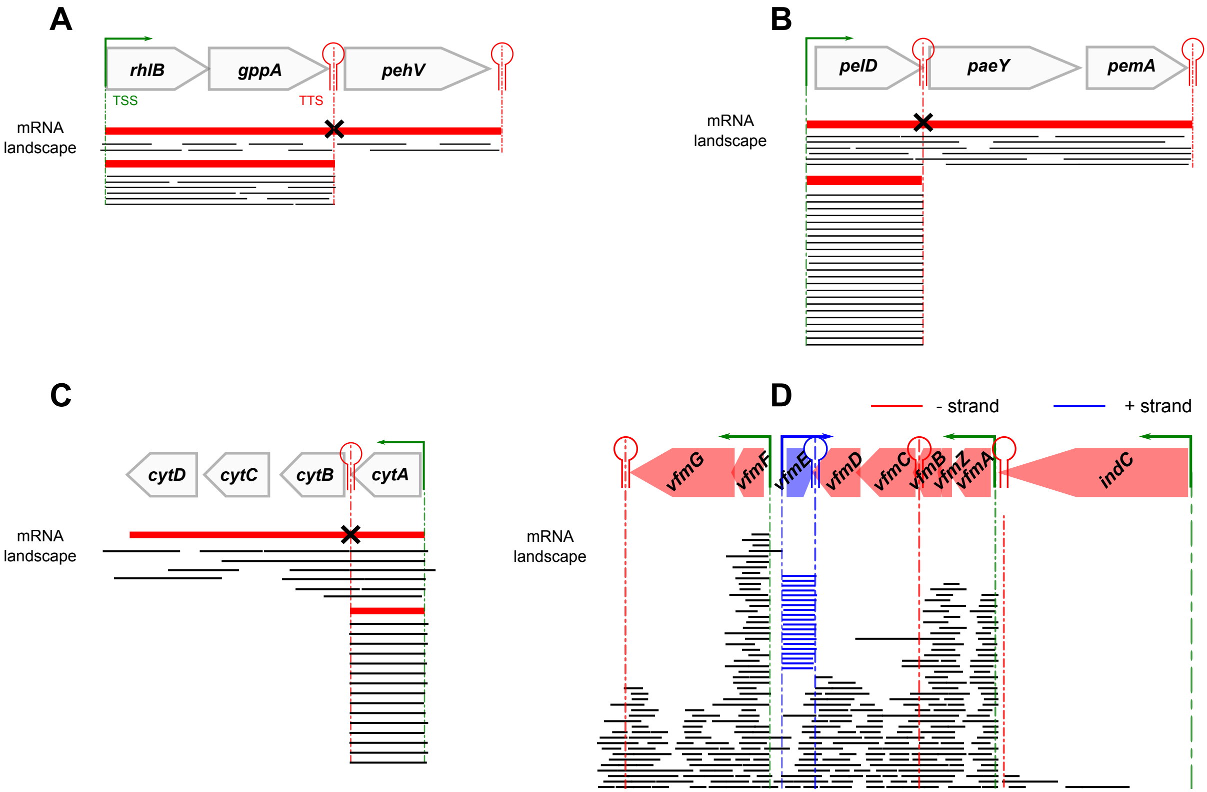

### Supplementary Fig. S3

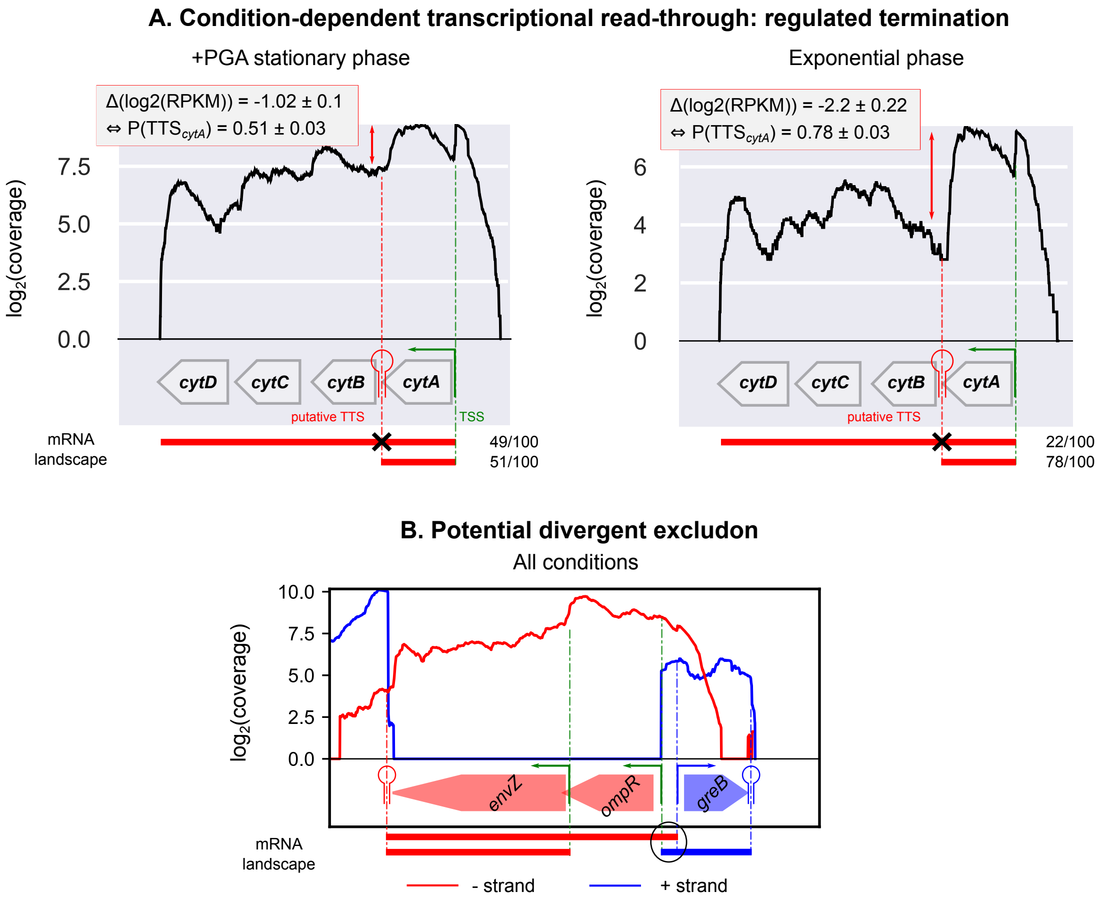

### Supplementary Fig. S4

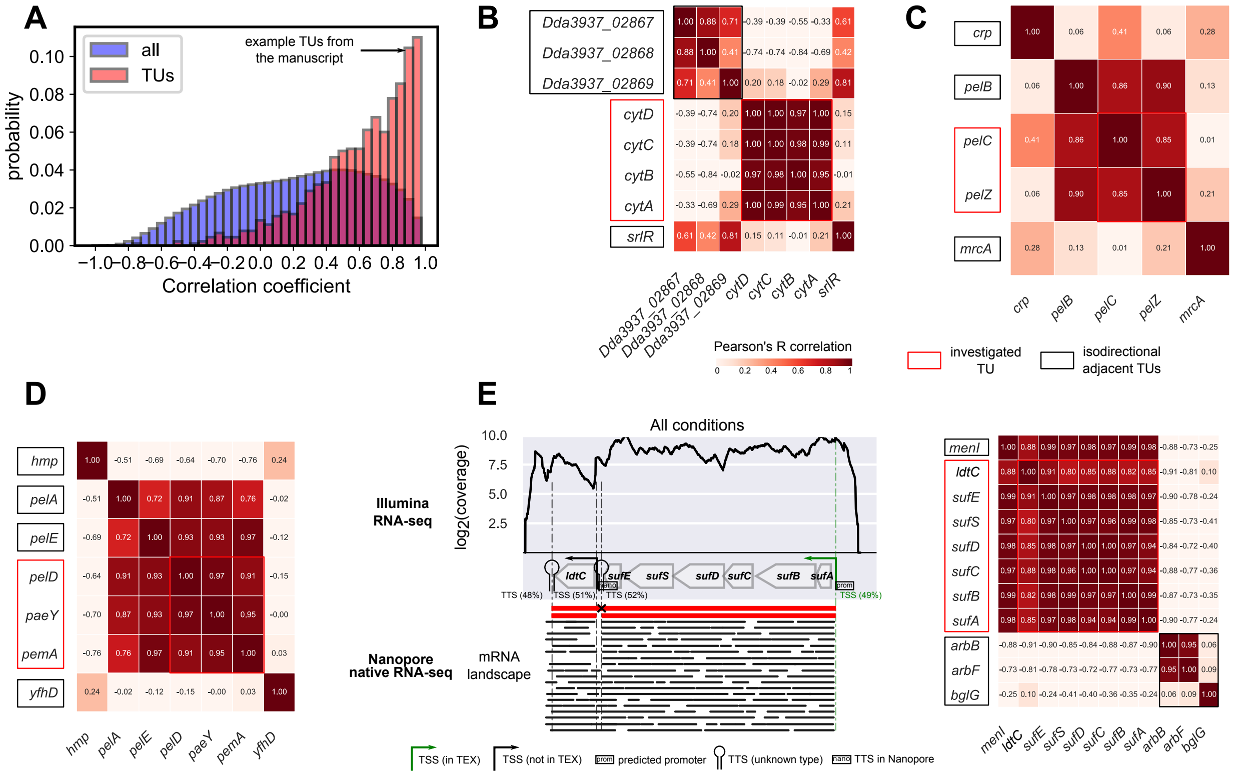

### Supplementary Fig. S5

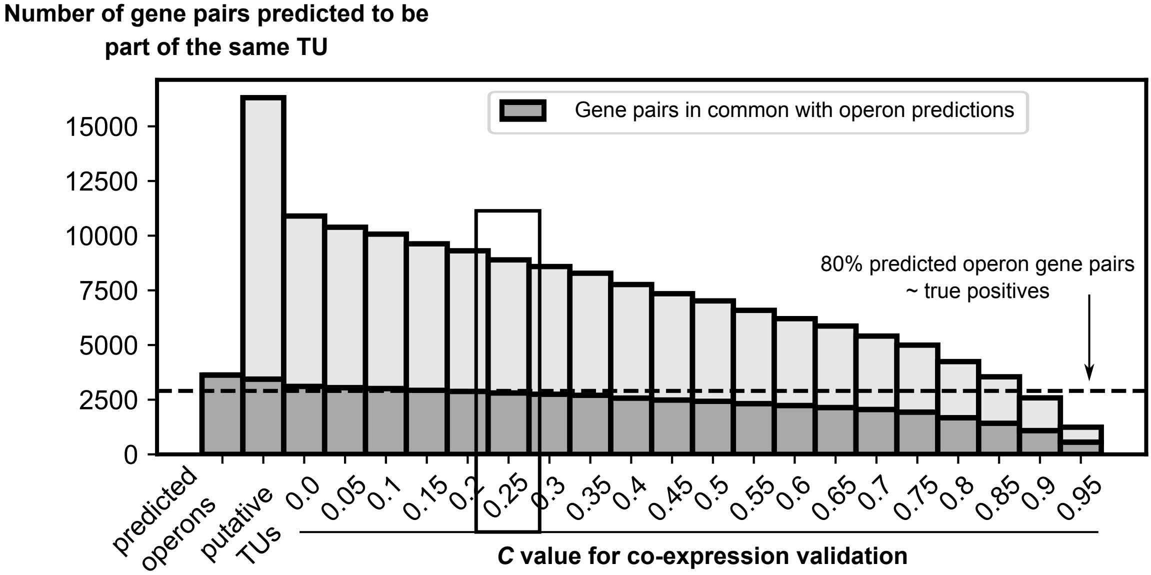
